# Pareto Suboptimal Resource Allocation and Microbial Growth

**DOI:** 10.64898/2026.09.08.749853

**Authors:** Valentina Baldazzi, Francis Mairet, Tomas Gedeon, Hidde de Jong

## Abstract

Microbial growth has often been analyzed under the assumption that microorganisms have evolved to optimize phenotypic characteristics of interest, such as growth rate and growth yield. This assumption has been useful, for example, for the genome-scale modeling of metabolism and the study of the allocation of cellular resources to physiological processes. In many experimental situations of interest, however, microorganisms are found to be suboptimal with respect to phenotypic characteristics that are thought to be favorable in that situation. Whereas a strong theoretical framework exists to mathematically relate cellular resource allocation strategies to Pareto optimality of microbial growth and other biological processes, much less is known about the consequences of Pareto suboptimality. We extend the framework to the latter case and show that a given Pareto suboptimal phenotype can be explained by a range of underlying resource allocation strategies, each corresponding to a different growth physiology and biomass composition. We test the predictions with the help of a coarse-grained model of microbial growth and published experimental data, which relate Pareto suboptimal rate-yield phenotypes of *Escherichia coli* and the microalga *Tisochrysis lutea* to the macromolecular composition of the cells (storage metabolite and total protein contents). Changing the focus from Pareto optimality to suboptimality provides interesting leads to exploring the diversity of growth strategies that can support a given phenotype. This change of perspective is of practical interest, because some of the growth physiologies within this range may be important for biotechnological applications.

**Author summary:** Many methods for the analysis of metabolism and growth are based on the assumption that, under the pressure of natural selection, microorganisms have evolved towards phenotypes that are optimal with respect to ecologically important characteristics. In mathematical terms, this assumption can be formulated as Pareto optimality of microbial growth and related to underlying resource allocation strategies that shape cellular physiology. Based on the observation that this assumption is often not satisfied in practice, we investigate the consequences of adopting the opposite assumption, namely that microbial growth and cellular resource allocation are suboptimal. We extend an existing mathematical framework to the case of Pareto suboptimality, which allows us to make predictions on the diversity of the macromolecular composition of microbial cells. These predictions are shown to correspond with experimental data on growth rate, growth yield, and macromolecular composition of the model bacterium *Escherichia coli* and the microalga *Tisochrysis lutea*. Our results thus associate suboptimality with observed metabolic diversity in microorganisms.

## Introduction

A commonly-made assumption in the analysis of microbial growth, and the analysis of biological systems more generally, is that organisms evolved to optimize a phenotypic characteristic or combination of characteristics [1, 2, 3, 4, 5, 6, 7]. Arguments supporting this optimality assumption are generally based on the mechanism of natural selection in evolution. The survival of a microbial species in its environment depends on the capacity to outcompete other species, that is, on the acquisition and conservation of values of phenotypic characteristics conferring a competitive advantage. Natural selection is therefore expected to favour microorganisms optimizing such characteristics.

Due to biophysical and biochemical constraints on cellular physiology, microorganisms cannot simultaneously optimize all phenotypic characteristics contributing to their fitness [8, 9, 10, 11]. That is, microbial cells need to make trade-offs between phenotypic characteristics competing for the same constrained cellular resources. Taking into account such conflicts between phenotypic characteristics, the optimality assumption can be formulated more precisely by saying that the functioning of organisms is expected to be Pareto optimal [8, 12]. A behavior is Pareto optimal when, among a set of phenotypic characteristics contributing to the fitness of the organism, increased performance on one characteristic comes at the price of decreased performance on another characteristic.

The optimality assumption sketched above has been fruitfully used in practice, in various forms, for the analysis of microbial growth. One example is Flux Balance Analysis (FBA), where a stoichiometric model of microbial metabolism is combined with constraints on the uptake of limiting substrates and other reactions to predict the maximum rate of biomass synthesis and the underlying metabolic flux distributions [13, 14]. FBA and its variants have been shown capable of quantitatively predicting the growth of microorganisms in different scenarios [15, 16, 17, 18]. Another example are studies of cellular resource allocation, that is, the relation between cellular investment in different classes of proteins and microbial growth. Assuming that cells allocate their resources so as to maximize their growth rate, coarse-grained models of microbial growth can reproduce the so-called ribosomal growth law, predicting a linear relationship between growth rate and ribosome content [19, 20, 21, 22, 23, 24].

A major argument against the use of optimality in these examples is that in many commonly studied environments, microorganisms are not found to be optimal with respect to phenotypic characteristics thought to be favorable in that environment. For example, when rapidly shifting *E. coli* cells from minimal to rich medium, ribosome activity increases on a time-scale faster than that required for the synthesis of new ribosomes, suggesting the presence of an unused ribosome reserve before the upshift [25, 26]. This is not consistent with the assumption, used in the above-mentioned derivation of the ribosomal growth law, that cells allocate resources to ribosomes and other proteins so as to maximize their growth rate. The unused ribosome reserve could have been exploited instead to further increase the growth rate. Another example is provided by the results of laboratory evolution experiments in which *E. coli* cells were propagated over tens of thousands of generations in a standard batch experiment [27]. When analyzing the measured growth rates and growth yields of both parent and evolved strains, there is little evidence that the bacteria are Pareto optimal with respect to these characteristics [28]. This agrees with the conclusions from a study in which hundreds of rate-yield phenotypes were compared with a theoretically predicted Pareto frontier [29] (see also [30]).

The above counterexamples are puzzling, but not sufficient to invalidate the optimality assumption. First, it is possible that microorganisms optimize phenotypic characteristics different from those that we intuitively consider important. For example, it was found from metabolic flux data that a variety of microbial strains are close to the Pareto frontier located in a three-dimensional space spanned by maximum ATP yield, maximum biomass yield, and the minimum sum of absolute fluxes [31]. However, the microorganisms are not Pareto optimal when the analysis is limited to only two of the three phenotypic characteristics. Second, the experimental conditions in the lab do not usually correspond to real-life conditions encountered by microorganisms. If evolution has adapted the physiology of microorganisms to a specific environmental niche, there is no *a-priori* reason that growth phenotypes are optimal in laboratory conditions. For example, while most experimental data come from controlled laboratory experiments where microorganisms are in a state of balanced growth with a single limiting nutrient, in real life microorganisms are submitted to conditions in which nutrient availability is diverse and constantly fluctuating [32, 33, 34, 35].

It is not our intention to engage in a debate on the validity of the optimality assumption. However, it is safe to conclude that, in common laboratory experiments, microbial growth is often not Pareto optimal with respect to commonly considered phenotypic characteristics. While much work has gone into the analysis of (Pareto) optimality in microbial growth, as mentioned above, much less is known about the consequences of making the assumption that microbial growth is suboptimal. Here we address this question, both on the theoretical level inspired by mathematical optimization theory [12] and in the practical context of modeling microbial growth.

A first result from our analysis is that the same Pareto suboptimal phenotype can be attained for very different underlying physiological states, that is, states in which cells function very differently due to differences in the underlying resource allocation strategies. We explore the practical consequences of this result for the predicted growth rate and growth yield of *E. coli*. Second, whereas some suboptimal growth physiologies correspond to usually observed biomass compositions, others are characterized by unexpectedly low protein and high storage metabolite concentrations. We combine the predicted diversity of physiological states with experimental data on growth rate, growth yield, and concentrations of protein and storage metabolites for *E. coli* and algae [36, 37, 38, 39].

This shows that changing the focus from Pareto optimality to suboptimality provides interesting leads for exploring the range of possible growth physiologies in microorganisms. Increased emphasis on diversity of growth strategies supporting the same phenotype is also of practical interest, because some of the physiologies within this range may be important for biotechnological applications.

## Results

### Analysis of Pareto optimal phenotypes

A phenotypic characteristic is defined as an observable, measurable characteristic of a microorganism, for example the growth rate (in units h^−1^) or growth yield (in units gram dry weight of biomass produced per mol of substrate taken up). A specific value of a phenotypic characteristic is called, following established terminology in genetics, a phenotypic trait, for example a growth rate of 0.2 h^−1^. The ensemble of relevant phenotypic traits is referred to as the phenotype. A phenotypic trait is determined by the physiology of a microorganism, *i*.*e*., the different processes taking place inside the cell, in interaction with the environment.

As explained above, in order to survive in a changing environment, microorganisms are assumed to simultaneously optimize multiple, possibly conflicting phenotypic characteristics. This type of problem is often encountered in economic theory, where the mathematical framework of Pareto optimization was developed, but it is also ubiquitous in biology. Phenotypic characteristics to optimize are usually called objectives in optimization theory and the problem of optimizing multiple, possibly conflicting objectives, a multiobjective optimization problem [12]. We introduce the key concepts in this section by means of an example with two phenotypic characteristics. The precise and fully general definitions and results, some of which summarize previous work [8, 9, 40], can be found in the *Methods* section.

Consider the problem of simultaneous maximization of growth rate and growth yield by a microorganism. On the physiological level, the limiting resource for microbial growth is usually considered to be proteins, the main component of cellular biomass. Proteins function as enzymes in carbon and energy metabolism and they also constitute the molecular machines responsible for the synthesis of macromolecules, in particular proteins themselves. Different allocations of the proteome to these different classes of protein, that is, different allocation strategies, will yield different growth rates and growth yields (see [2, 41, 42] for reviews).

Mathematically, each allocation strategy will be a vector of allocation variables *x ∈*ℝ^*n*^, where *x* corresponds to the proteome fractions assigned to *n* different processes. The set *X* of all possible allocation strategies can be described by a set of constraints of the form *g*_1_ (*x) ≤* 0, *…, g*_*p*_ *(x) ≤* 0, and is called feasible set. Among all these strategies the cell is simultaneously trying to maximize growth rate *f*_1_(*x*)and growth yield *f*_2_(*x*):

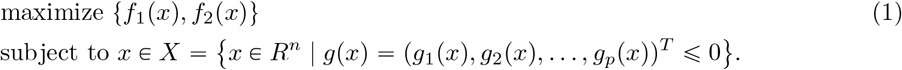

The problem defined by Eq. 1 is a multiobjective optimization problem with objective functions *f*_1_(*x*) and *f*_2_(*x*). We illustrate this problem in Figure 1 with two allocation variables *x*_1_, *x*_2_ and simple linear constraints *x*_1_ ≥ 0, *x*_2_ ≥ 0, and *x*_1_ *x*_2_ −1 ≤ 0, indicating that the proteome fractions are non-zero and sum to a maximum of 1. In Figure 1 the feasible set of resource allocation strategies *X* is pictured on the left. The set of corresponding phenotypic traits *P, i*.*e*., all potential values of phenotypic characteristics ( *f*_1_ (*x), f*_2_ (*x)), x ∈ X*, is shown on the right. The set *P* is the set of feasible phenotypes.

**Figure 1.**
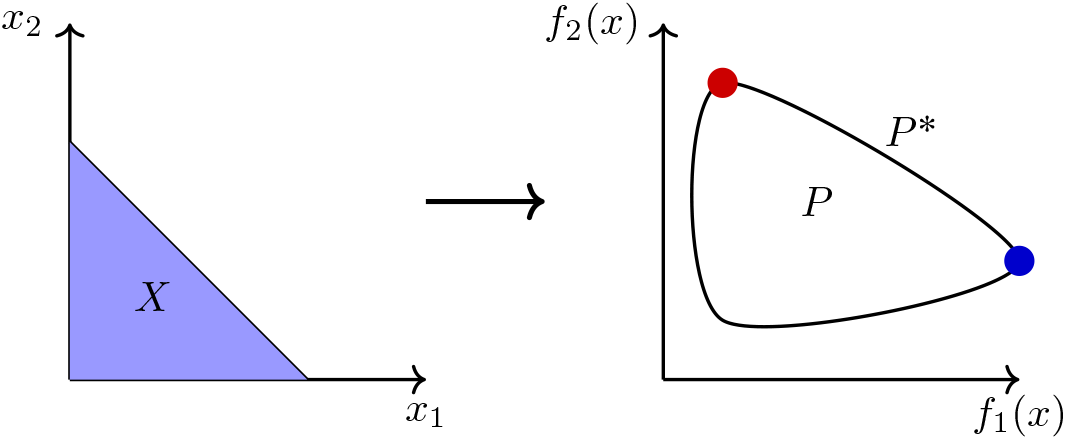
Mapping from set of resource allocation strategies to set of phenotypes and Pareto frontier. The pair of objective functions *f =* (*f*_1_, *f*_2_) maps a domain *X* to a closed and bounded set *P* . The objective functions are phenotypic characteristics like growth rate and growth yield, which are functions of the underlying resource allocation strategy of the cell. The Pareto frontier *P* ^*^ is the part of the boundary where *f*_1_ and *f*_2_ cannot increase simultaneously. In the case of two objective functions, *P* ^*^ is a one-dimensional curve segment that joins the maximum of *f*_1_ (blue) to the maximum of *f*_2_ (red).

We say that a pair of traits (*p*_1_, *p*_2_) dominates another pair 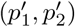 if 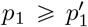 and 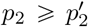, and at least one of the inequalities is strict. In this case, the phenotype (*p*_1_, *p*_2)_ is preferred to 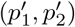 because at least one of the values for the characteristics to be maximized is higher in the first phenotype. The set of traits (*p*_1_, *p*_2_) that is not dominated by any other feasible 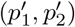, that is, the set of points in *P* where one cannot simultaneously improve both phenotypic characteristics *f*_1_ and *f*_2_, is called the Pareto frontier *P* ^*^. Since along the Pareto frontier the increase in one phenotypic characteristic forces a decrease in the other, the entire set *P* ^*^ consists of solutions of the multiobjective optimization problem of Eq. 1. These solutions are called Pareto optimal phenotypes and the corresponding set of all allocation variables *x ∈ X* that lead to Pareto optimal phenotypes is called the Pareto set.

Previous work has related the existence and the form of a Pareto frontier *P* ^*^ to properties of both the functions *f* relating resource allocation strategies to phenotypic characteristics and the space of possible strategies *X* [8, 9, 40].

### Theorem 1 (**Pareto optimality**). Provided that

Assumption 1: the objective functions *f*_1_, *f*_2_ and the constraint functions *g*_*i*_ are continuously differentiable,

Assumption 2: each function *f*_1_, *f*_2_ has a unique maximum in the interior of the feasible set *X*, obtained for strategies *m*_1_, *m*_2_,

Assumption 3: the maxima are different, *i*.*e*., *m*_1_ ≠ *m*_2_,

Assumption 4: the functions *f*_1_, *f*_2_ and the set *X* are strongly convex, it holds that

1. there exists a Pareto frontier *P* ^*^,
2. the Pareto frontier *P* ^*^ is a curve that joins the maximum of *f*_1_ to the maximum of *f*_2_,
3. the multiobjective optimization problem of Eq. 1 can be formulated as a collection of single objective optimization problems

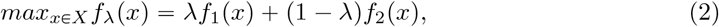

where 0 ≤ *λ ≤* 1 indicates the relative preference for *f*_1_ and *f*_2_. For well-behaved problems that satisfy the above assumptions, the Pareto frontier *P* ^*^ will be exactly the set of optima for Eq. 2 as the value of *λ* changes between 0 and 1.

The example in Figure 1 has been chosen such that it satisfies the above property. As can be seen, the Pareto frontier runs from the maximum growth rate *f*_1_ (*λ* = 1) to the maximum growth yield *f*_2_ (*λ =* 0). The reformulation of the optimization problem to Eq. 2 emphasizes that at each point on the Pareto frontier the cell maximizes a particular mixture of growth rate and growth yield.

While Assumptions 1 and 3 are generic, in the sense that they only exclude edge cases, Assumptions 2 and 4 meaningfully restrict the set of functions that we allow. The intuition of the role played by the latter two assumptions can be conveyed by sketching the level curves of the functions *f*_1_ and *f*_2_ in the feasible set of resource allocation strategies *X* (Figure 2). Consider a point in *X*

**Figure 2.**
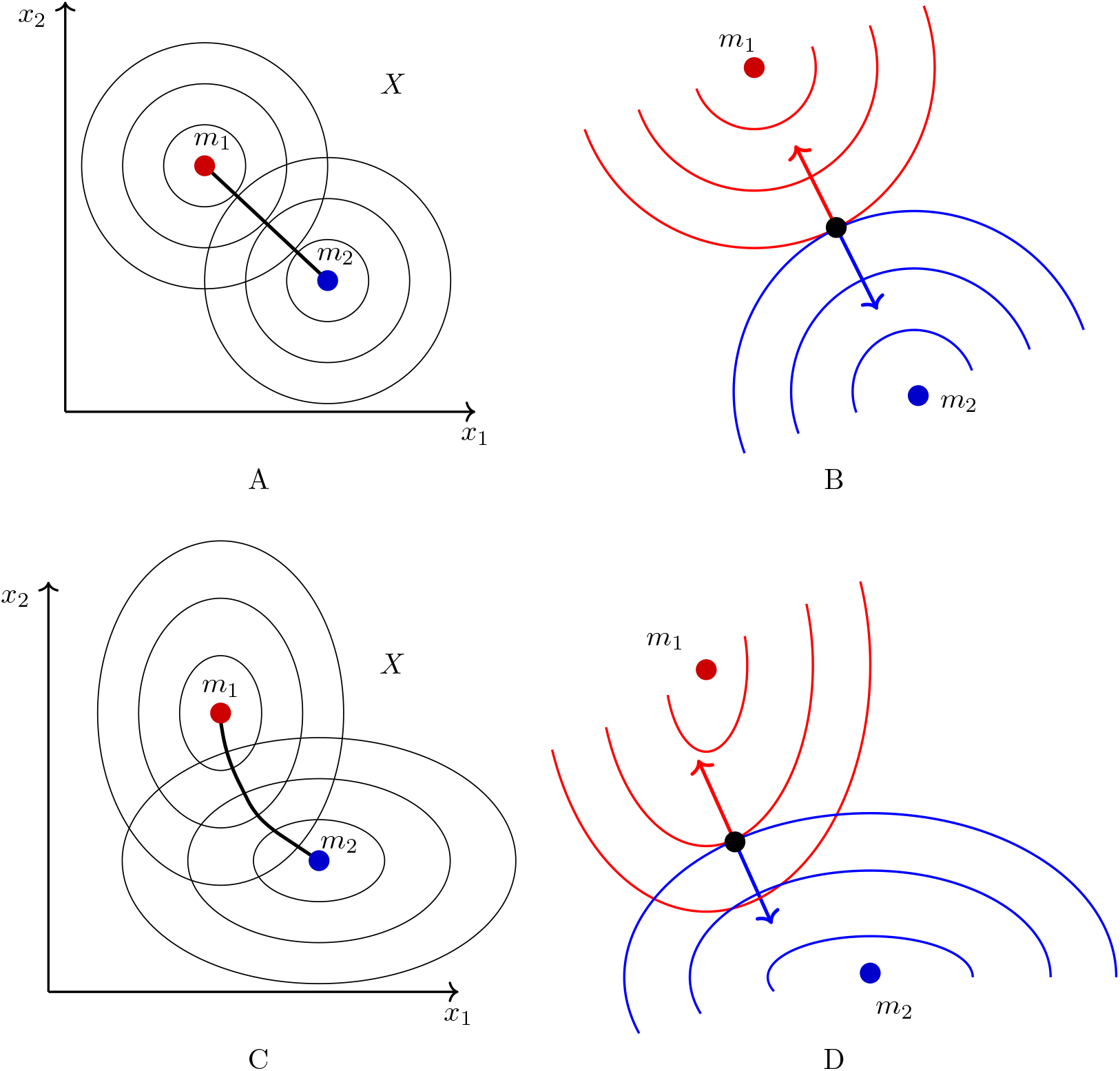
Geometrical representation of the Pareto frontier in the feasible set of resource allocation strategies. (A) Pareto set in the domain *X*. The point *m*_1_ (*m*_2_) is the point at which the maximum of function *f*_2_ (*f*_1_) is attained. We assume that these points are in the interior of the domain *X*. If the level curves are circles, the Pareto set is the black line segment connecting *m*_1_ and *m*_2_. (B) A point corresponding to a resource allocation strategy is included in the Pareto set when it holds at this point that *∇ f*_1_ = *ν∇ f*_2_ for some negative *ν*. (C)-(D) When the level curves are not circles but ellipses, the Pareto set may no longer be a line segment, but convexity of *f*_1_ and *f*_2_ still guarantees that the set is a one-dimensional curve segment.

where the direction of increase in *f*_1_ is exactly opposite to the direction of increase of *f*_2_. In fact, this is the defining (local) property of a Pareto set of resource allocation strategies: a Pareto set is the set of points where *∇ f*_1_ = *ν∇ f*_2_ for some negative *ν* [12]. At these points, the level curves of *f*_1_ and *f*_2_ are tangent, but the gradients of *f*_1_ and *f*_2_ point in opposite directions. If the level curves are circles, the Pareto set is the line segment that connects points *m*_2_ (the unique maximum of *f*_2_) and *m*_1_ (the unique maximum of *f*_1_), see Figure 2A-B [8]. When the level curves are not circles, but general convex curves encircling their respective maxima, the gradient vectors may not always point directly at *m*_1_ and *m*_2_. As a consequence, the Pareto set may not be a line segment, but the convexity of *f*_1_ and *f*_2_ still guarantees that the set is a one-dimensional curve segment [40, 9], see Figure 2C-D.

We consider Assumptions 2 and 4 biologically reasonable, as confirmed in the example discussed in detail below. However, the results summarized above can be generalized in case the assumptions are not satisfied, that is, when *f*_1_ and *f*_2_ have multiple maxima and/or are not convex [40, 9]. This may lead to situations in which the Pareto set of resource allocation strategies and/or the Pareto frontier in the phenotype space are no longer continuous. We will disregard these situations here.

### Analysis of Pareto suboptimal phenotypes

In spite of the significant theoretical appeal of the idea that microorganisms operate close to a Pareto frontier, in many situations of practical interest, this does not seem to be the case. For example, the *Introduction* mentioned that there is not much experimental evidence confirming that microorganisms optimize common phenotypic characteristics like growth rate, growth yield, or a combination of the two in standard laboratory conditions. This discrepancy could be explained by assuming that microorganisms are not optimally adapted to the environment in which they are studied or that they do not actually optimize the chosen combination of phenotypic characteristics.

Figure 3 schematically develops these explanations for non-optimality by projecting data points on the set of predicted feasible phenotypes *P* . Panel A shows that the observed phenotypes fall below the Pareto frontier, assuming that microorganisms optimize *f*_1_ and *f*_2_. The microorganisms are therefore Pareto suboptimal,. However, our assumption that the microorganisms optimize *f*_1_ and *f*_2_ could be misguided. In particular, for another combination of phenotypic characteristics, say *f*_1_ and *f*_3_, the same observations could lie close to the new Pareto frontier, as shown in panel B. Alternatively, if, in addition to *f*_1_ and *f*_2_ we also consider *f*_4_, the points located far from the Pareto frontier in panel A may end up close to the new Pareto frontier in panel C, which is a two-dimensional surface in a three-dimensional phenotype space. This example illustrates that, when considering a subset of phenotypic characteristics, we are not guaranteed to preserve Pareto optimality observed for the entire set.

**Figure 3.**
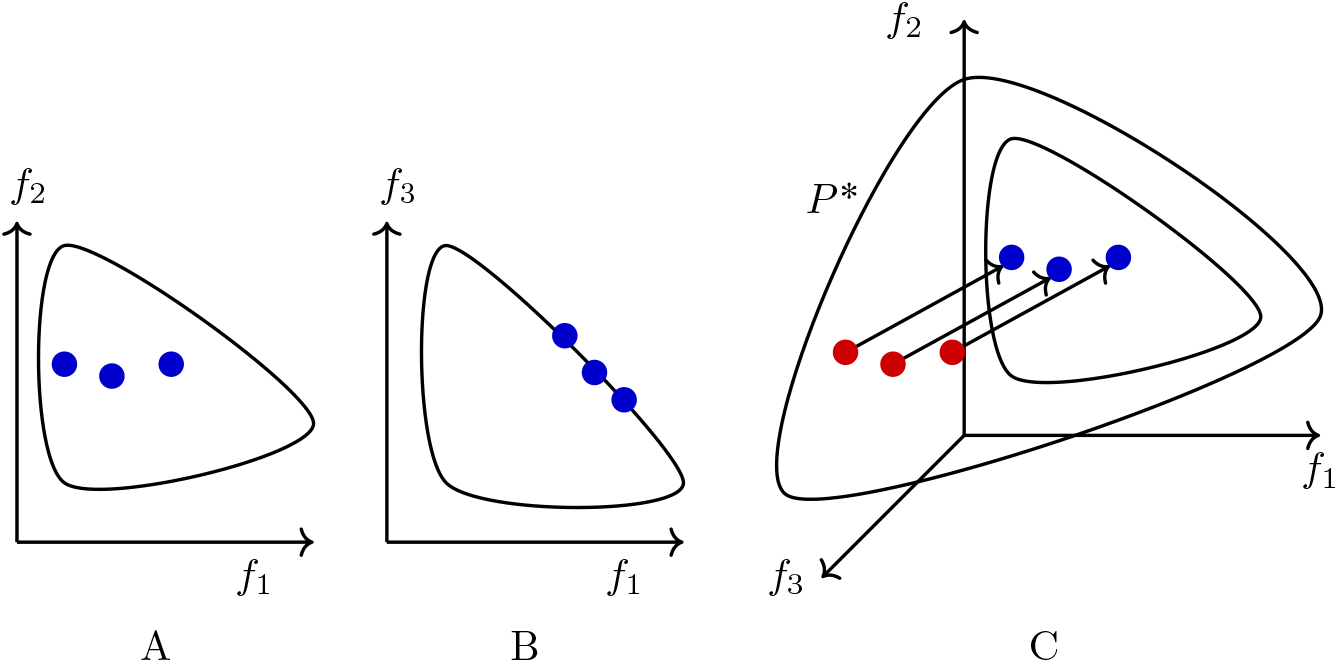
Schematic explanations of the occurrence of Pareto suboptimality. (A) Three data points (blue dots) are projected on a phenotypic space defined by the characteristics {*f*_1_, *f*_2_}. The observed phenotypes fall into the predicted feasible set, but they are located far from the Pareto frontier and therefore display Pareto suboptimality. (B) When the data points are projected on an alternative phenotypic space, defined by {*f*_1_, *f*_3_}, they are located close to the Pareto frontier and therefore display Pareto optimality. (C) Consider a three-dimensional phenotypic space defined by {*f*_1_, *f*_2_, *f*_3_}, where the Pareto frontier *P* ^*^ is a two-dimensional surface. Whereas the data points (red) lie on the Pareto frontier for the full set of characteristics, they show Pareto suboptimality on the strict subset of panel A. Arrows indicate the projection of the data points to the two-dimensional subspace of characteristics {*f*_1_, *f*_2_} ⊂ {*f*_1_, *f*_2_, *f*_3_}.

In practice, it is difficult to decide whether Pareto suboptimality occurs because microorganisms are not optimally adapted to their environment in which they evolve or because we project their phenotypes to a combination of characteristics that does not adequately account for the structure of that environment. For commonly considered characteristics, however, as discussed in the *Introduction*, one often finds that the observed phenotypes are located away from the predicted Pareto frontier, that is, microorganisms have Pareto suboptimal phenotypes. This makes it interesting to investigate the consequences of Pareto suboptimality.

Following the logic that led to Theorem 1, a general result on Pareto suboptimality can be formulated. Like in the previous section, we consider the case of two phenotypic characteristics, *e*.*g*., growth rate and growth yield, defined by the objective functions *f*_1_ and *f*_2_, respectively.

However, the results can be generalized to more than two characteristics (*Methods*):

#### Proposition 2

(**Pareto suboptimality**). Retain Assumptions 1-4 from Theorem 1 and make the further assumption that the Pareto set lies in the interior of *X*. Then, for any feasible phenotype dominated by the Pareto frontier *P* ^*^, *p* ∈ *P* \*P* ^*^, and lying in the neighborhood of *P* ^*^, there exist feasible *x, x*^’^ ∈ *X, x* ≠ *x*^’^, such that (*f*_1_(*x*), *f*_2_(*x*)) “(*f*_1_(*x*^’^), *f*_2_(*x*^’^)) = *p*.

In other words, a given suboptimal Pareto phenotype can be obtained for several different, feasible underlying resource allocation strategies. These strategies will be called equivalent. The proposition is illustrated in Figure 4A, for the same example as in Figure 2. Consider again level curves for *f*_1_ and *f*_2_ in *X*, with maxima obtained for *m*_1_ and *m*_2_. Due to the convexity of *f*_1_ and *f*_2_ (Assumption 1), the level curves are convex. Any given pair of level curves for *f*_1_ and *f*_2_ therefore has either no intersection, a single intersection, or two intersections. The case of no intersection is not of interest here, because it does not lead to a feasible phenotype. When two level curves touch, and thus have a single intersection, the corresponding phenotype is included in the Pareto frontier (Figure 2). When the level curves intersect twice, the resource allocation strategies corresponding to the intersections are equivalent, in the sense that they map to identical phenotypes. But this phenotype is suboptimal, in the sense that it is located away from the Pareto frontier *P* ^*^.

**Figure 4.**
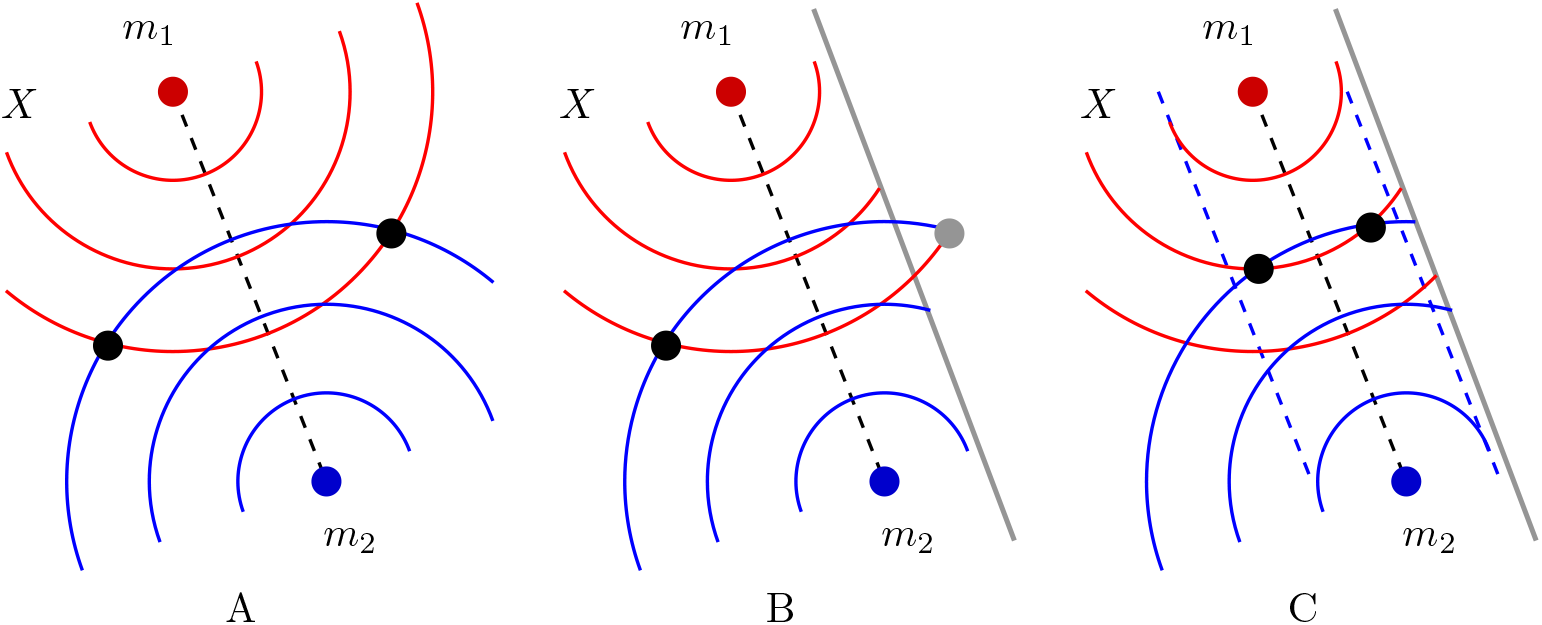
Geometrical representation of resource allocation strategies that give rise to Pareto suboptimal phenotypes. (A) If convex level curves of *f*_1_ and *f*_2_ contained in the feasible set *X* intersect each other, they do so in two points (black), while they intersect each other in a single point along the Pareto frontier (dashed curve). (B) If the convex level curves are not entirely contained in *X*, some of the intersections may not be feasible. The vertical line denotes the boundary of *X* and the gray point the intersection of two level curves outside *X*. (C) If the Pareto frontier is in the interior of the feasible region, there is always a region surrounding it (blue dashed curves), where the level curves intersect in two feasible points.

Proposition 2 requires suboptimal phenotype *p* to be located in the neighborhood of Pareto frontier *P* ^*^ and that the Pareto set lies in the interior of *X*. In the example of Figure 4, we illustrate why this must be the case. The level curves of *f*_1_ and *f*_2_ in *X ⊂* ℝ^2^ are one-dimensional curves, which generally intersect in two points *x, x*^’^, as shown in panel A. When the level curves are not entirely included in *X*, however, one of the resource allocation strategies may not be feasible, as shown in panel B. This situation can be avoided if *p* is sufficiently close to *P* ^*^ and *P* ^*^ is in the interior of *X*, since then both intersections are guaranteed to lie within *X* (panel C). The restriction that *p* is close to *P* ^*^ is made solely to facilitate the intuitive discussion of Figure 4. A general result in the *Methods* section shows that for a problem with *n ≥* 2 resource allocation strategies, there is a dense set of values *p∈ P* for which the set of equivalent strategies producing phenotype *p* is of dimension *n −* 2. It follows that for *n >* 2, the set of equivalent strategies is infinite.

Equivalent strategies have the same phenotype, but correspond to different allocations of cellular resources to proteins, and therefore to cells with a different physiology. If the equivalent strategies are located close to each other, in the neighborhood of the Pareto set (Figure 4), then one expects the corresponding physiological states to be close as well. For equivalent strategies lying farther apart, the physiological states giving rise to the same phenotype may be quite distinct. How can microorganisms with very different states of the underlying metabolic and gene expression processes nevertheless exhibit the same phenotype? In the next section, we consider these questions in the context of a concrete example, a coarse-grained model of bacterial growth.

### Pareto suboptimal growth of microorganisms: prediction of alternative physiological states

In recent years, a large number of resource allocation models of microbial growth in the spirit of those described in the previous sections have appeared (see [2, 41, 42, 43] for reviews). Some of these models are best conceived as extensions of large-scale, constraint-based models of microbial metabolism [30, 44, 45, 46, 47, 48], completing the metabolic network of the cell with reactions for the synthesis of proteins and other macromolecules. This leads to additional constraints on the reaction rates due to the allocation of limiting protein resources to specific transporters and enzymes, as well as ribosomes, catalyzing these reactions. Other models can be seen as coarse-grained representations of the cellular processes involved in microbial growth, reducing the complex reaction network to a limited number of macroreactions responsible for major growth-related processes [19, 20, 22, 23, 24, 49, 50, 51, 52, 53, 54, 55]. In the latter models, resource allocation is simplified to the partitioning of the proteome into different protein categories, each of which is responsible for a specific macroreaction. Despite these simplifications, coarse-grained models have been shown capable of accounting for major aspects of microbial growth phenotypes, such as the ribosomal growth law and the occurrence of a switch from respiration to fermentation.

Although there are differences between the different types of coarse-grained model, which determine the possible analyses that can be done and the conclusions that can be drawn from these analyses, they all support the optimization analysis discussed here: find a resource allocation strategy optimizing a specific phenotypic characteristic, such as growth rate or growth yield. In this work, we especially consider coarse-grained models of microbial growth. A first reason is that the smaller size of such models allows for a more straightforward comparison of the physiological states underlying Pareto optimal and Pareto suboptimal growth. A second reason is more technical. Many (though not all) large-scale models of microbial growth are based on flux constraints and do not provide kinetic expressions for the reaction rates. As a consequence, the underlying system of equations is degenerate and does not generally predict a single phenotype or a single combination of phenotypes for a given (optimal or suboptimal) resource allocation strategy. The fact that the mapping from *X* to *P* is not a function precludes the application of the theoretical results presented in the previous sections.

In what follows, we focus on a specific coarse-grained model providing a combined description of the carbon and energy balances in microbial growth [29] (Figure 5). The model includes macroreactions for substrate uptake and catabolism, the generation of ATP by respiratory and fermentative pathways, as well as the synthesis of proteins and other macromolecules. For our present purpose, we extended the model with a macroreaction for the synthesis of glycogen, a storage compound that is a homopolymer of glucose [56] (*Methods*). The proteome is divided into five classes of proteins: ribosomes and other translation-affiliated proteins, enzymes in central carbon metabolism, enzymes in respiration and fermentation metabolism, and a residual category of other proteins responsible, among other things, for the synthesis of other macromolecules, notably RNA and DNA. Each macroreaction in the model is catalyzed by one class of proteins. The fermentative and respirative pathways produce ATP, which is (mostly) consumed by the synthesis of macromolecules. Fermentation leads to the incomplete catabolization of the substrate, here assumed to be glucose, and the secretion of a by-product, here acetate. The model also accounts for the turnover of macromolecules, responsible for a large fraction of cellular maintenance costs, as well as energy dissipation due to ATP spillage. One particularity of the model compared to other models of its type is that biomass not only includes proteins but also other cellular components, in particular other macromolecules and metabolites. The macroreactions are described by kinetic equations. The parameters in these equations were determined, without parameter fitting, from prior experimental data for the *E. coli* strain BW25113 growing in batch in minimal medium with glucose (see *Text S1* for details).

**Figure 5.**
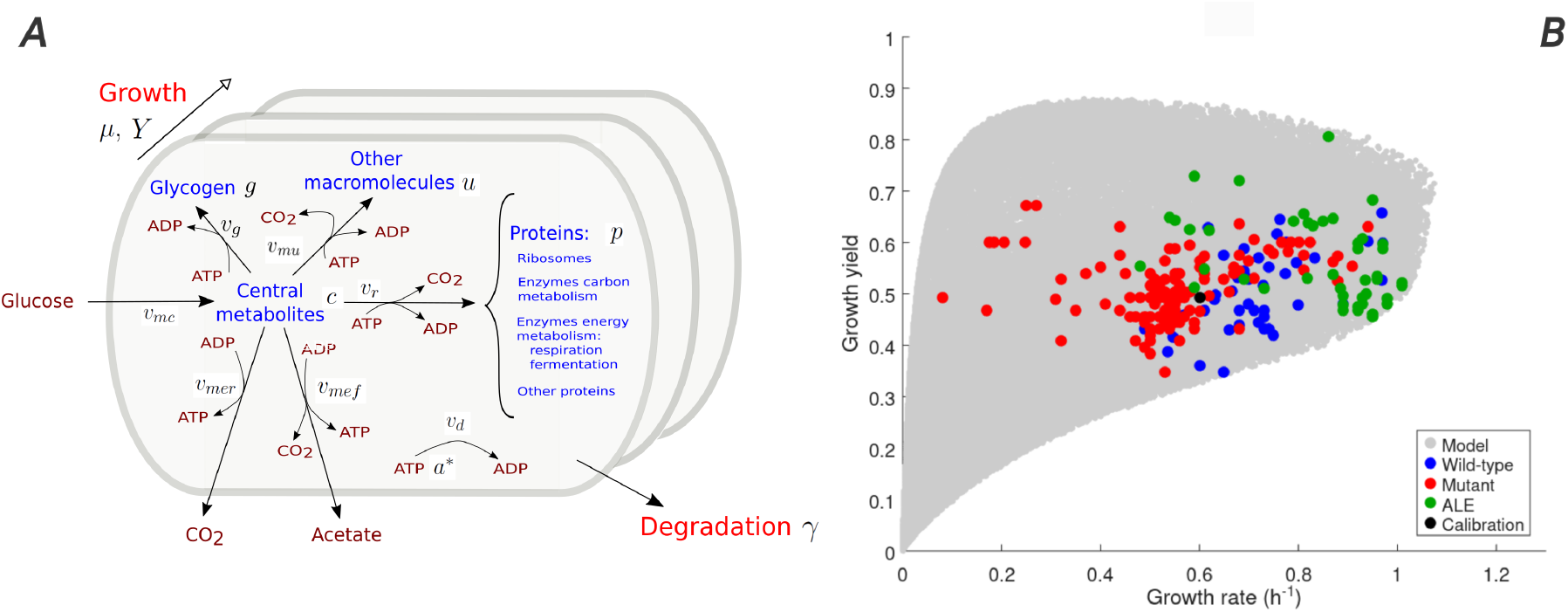
Coarse-grained model of microbial growth with coupled carbon and energy balances. (A) Schematic outline of the model, showing biomass constituents and macroreactions, as well as growth and degradation of biomass. (B) Predicted and observed rate-yield phenotypes during balanced growth of *E. coli* on minimal medium with glucose. Each feasible resource allocation strategy leads to a predicted phenotype in the rate-yield plane (grey dot). The observed rate-yield phenotypes concern the BW25113 strain used for calibration of the model, other wild-type strains, mutant strains obtained by directed mutagenesis, and mutant strains from adaptive laboratory evolution (ALE) experiments (black, blue, red, and green dots, respectively). The model originates from previous work [29], but has been extended with a macroreaction for the synthesis of storage compounds, in particular glycogen (*Methods*).

The prediction of the phenotypic characteristics of interest here, growth yield and growth rate, requires numerical solution of a system of kinetic equations at steady state, knowing that a resource allocation strategy always maps to a single steady state [29]. Figure 5B shows the set of rate-yield phenotypes (grey region) resulting from uniform sampling of the space of feasible resource allocation strategies. The growth rate reaches a maximum of around 1.1 h^−1^. This maximum ultimately depends on the biomass concentration, assumed constant in the model, which puts an upper bound on the enzyme and ribosome concentrations required to take up and assimilate glucose from the environment. The growth yield is expressed as mmol of carbon included in the biomass per mmol of carbon taken up. As a consequence, the growth yield varies between 0 and 1. The feasible traits predicted from the model correspond rather well with a database of measured rates and yields reported in the literature (colored dots). The database includes the BW25113 strain used for model calibration, other common *E. coli* wild-type strains, strains with mutants in regulatory genes, and strains obtained from adaptive laboratory evolution (ALE) experiments [29].

It can be verified that the growth model satisfies the assumptions of Theorem 1. The set of feasible resource allocation strategies is determined by the constraints that the values of the individual resource allocation variables lie between 0 and 1, and that they sum to 1. The resource allocation vector is defined as *x* = (*x*_*r*_, *x*_*c*_, *x*_*er*_, *x*_*ef*_, *x*_*u*_), partitioning cellular protein resources over ribosomes and other translation-affiliated proteins (*x*_*r*_), enzymes in central carbon metabolism (*x*_*c*_), enzymes in respiration and fermentation metabolism (*x*_*er*_ and *x*_*ef*_ ), and a residual category of other proteins (*x*_*u*_), respectively. The resource allocation vector satisfies the following (in)equalities:

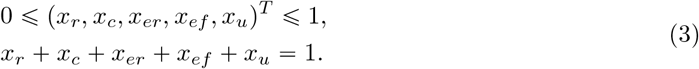

The feasible set *X* defined by these constraints is convex and the (in)equalities are continuously differentiable in the resource allocation variables. The functions *f*_1_ and *f*_2_ mapping a resource allocation strategy to a growth rate and a growth yield, respectively, are implicitly defined by the evaluation of these quantities at steady state. Under the assumption that *x*_*u*_ is constant, and given that *x*_*ef*_ can be expressed using Eq. 3 as a function of the other variables, *f*_1_ and *f*_2_ are functions of *x*_*r*_, *x*_*c*_, *x*_*er*_. When projecting the values of the set of feasible phenotypes *P* on the subspace of *X* defined by *x*_*r*_, *x*_*c*_, *x*_*er*_ (Figure 6A), we see that the functions *f*_1_ and *f*_2_ have a single maximum in *X* and that these two maxima are different. Moreover, the projection reveals that *f*_1_ and *f*_2_ are continuously differentiable and convex functions of the resource allocation variables.

**Figure 6.**
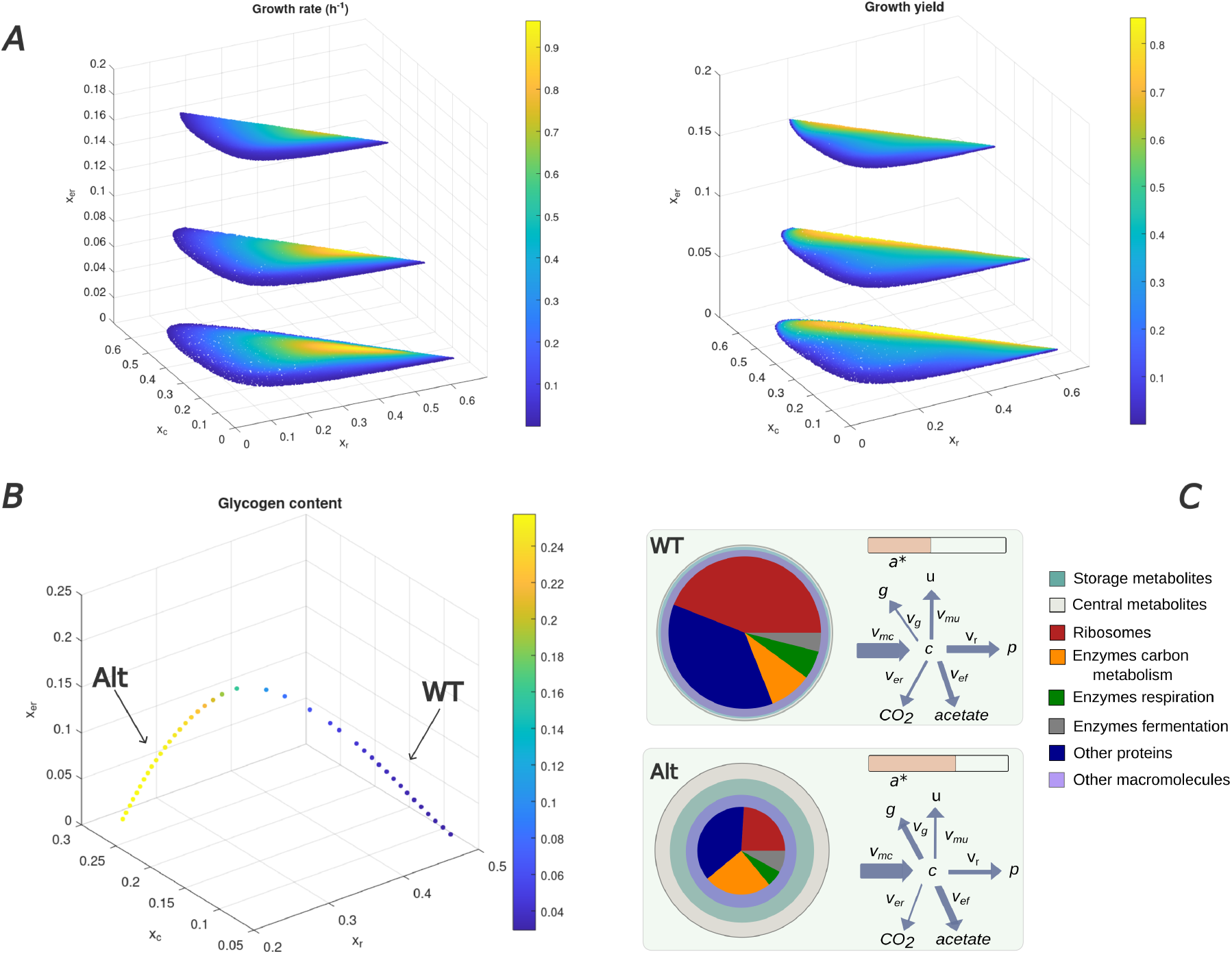
Theoretical analysis of Pareto suboptimal solutions of the coarse-grained model of microbial growth. (A) Predicted values of growth rate and growth yield obtained for feasible resource allocation strategies, in conditions of balanced growth on minimal medium with glucose. *x*_*u*_ was set to the constant value of 0.37 for the BW25113 reference strain, while three different values for *x*_*er*_ were explored (0.02, 0.1, 0.2). Note that each combination (*x*_*u*_, *x*_*r*_, *x*_*c*_, *x*_*er*_) fixes the remaining value of *x*_*ef*_ due to the constraint that the resource allocation parameters sum to 1. (B) Equivalent strategies predicted to result in the same rate-yield phenotype as the BW25113 strain (*µ* = 0.61 h^−1^, *Y* = 0.5, [60]). The color code indicates the glycogen concentration. The strategy marked by BW corresponds to the experimentally-determined strategy of the BW25113 strain [62], the strategy marked by Alt to a predicted alternative strategy with a high glycogen concentration. (C) Pictograms summarizing the growth physiologies corresponding to the resource allocation strategies in panel B.

Theorem 1 now allows us to draw the conclusion that the model predictions exhibit a Pareto frontier and that this Pareto frontier is a curve that connects the maximum growth yield to the maximum growth rate, as confirmed by Figure 5B. Interestingly, common laboratory strains, like the BW25113 strain, lie far below the Pareto frontier. Clearly, the model is simple and likely neglects constraints that play a role in shaping the feasible rate-yield phenotypes, and therefore the location of the Pareto frontier, such as the membrane space available for transporters and the protein complexes making up the electron transport chain [57, 58, 59]. However, the plot in Figure 5B shows that several, especially ALE strains, lie considerably closer to the Pareto frontier than BW25113 and other common laboratory strains. This strongly suggests that the latter have Pareto suboptimal rate-yield phenotypes. Another study, using a large kinetic model of *E. coli* central metabolism reached a similar conclusion [30].

This conclusion motivates a closer look at the resource allocation strategies supporting suboptimal Pareto phenotypes. In particular, we focus on resource allocation strategies that give the same growth rate and growth yield as the BW25113 strain, that is, a growth rate of 0.61 h^−1^ and a growth yield of 0.5 [60, 61]. Assuming that the entire Pareto set of resource allocation strategies lies in *X*, Proposition 2 and further results in the *Methods* predict that an infinite number of different resource allocation strategies give the same rate-yield phenotype as the BW25113 strain. Figure 6B plots these strategies by projecting the strategies on the subspace defined by *x*_*r*_, *x*_*c*_, *x*_*er*_, for different values of *x*_*ef*_ . The resource allocation strategy observed for the BW25113 strain [62], which was used for calibration of the model, is marked by WT for reference.

The different resource allocation strategies correspond to alternative physiological states predicted for *E. coli*. Figure 6C shows two examples of such states, one corresponding to that observed for the BW25113 strain and one for an alternative predicted state with a very different growth physiology. Whereas most of the biomass in the BW25113 strain growing on minimal medium with glucose consists of protein, the model indicates that the same growth rate and growth yield could be attained with a lower protein concentration. In the latter case, the biomass consists in large part of storage metabolites, notably glycogen. A major difference between the two strategies is that the bacteria have a much higher investment in ribosomes, in relative and absolute terms, in the first strategy. This allows a higher fraction of the central metabolites obtained from glucose to be converted into protein, leading to a higher total protein concentration. In the second strategy, the investment in ribosomes is much lower, with a correspondingly lower total protein concentration. The relative protein fraction of metabolic enzymes is higher in the second than in the first strategy though, endowing the bacteria with the capacity to take up glucose and transform it into (storage) metabolites. As a consequence, the mass fraction of glycogen is much higher in the second strategy. The glycogen concentration corresponding to the different alternative strategies is indicated in color code in Figure 6B. A higher glycogen concentration correlates with a lower total protein concentration (not shown).

The coarse-grained model of microbial growth considered in this section illustrates one of the main conclusions from the theoretical analysis of Pareto suboptimality in the previous section: the existence of multiple resource allocation strategies yielding the same phenotypic characteristics. When dissecting the underlying physiological states, in particular as regards their biomass composition, we see that the growth rate and growth yield of the strain used for calibration of the model are predicted to be feasible for quite different total protein concentrations. Whereas a high total protein concentration enables a high effective conversion rate of substrate to macromolecules, an alternative strategy would be to accumulate a large proportion of the substrate in the form of storage metabolites like glycogen. This conclusion directly follows from the logic of the model, but does there exist any experimental data in support? In the next two sections, we give some tentative answers to this question by exploiting data available in the literature.

### Experimental evidence for alternative physiological states in bacteria

Many model microorganisms used in the laboratory have a rather specific biomass composition. They mostly consist of protein, with a mass fraction varying, for *E. coli*, between 65% and 80% when the growth rate decreases from 1.3 to 0.3 h^−1^. The total mass fraction of metabolites, including storage metabolites such as glycogen, is usually minor, on average well below 10% [63].

The analysis of the model in the previous section revealed that the same growth rate and growth yield as that of a reference *E. coli* strain, grown in minimal medium with glucose, might be attained for a whole range of biomass compositions. Some of these are quite different from the typically measured values reported above (Figure 6C). Have bacterial strains with the same Pareto suboptimal phenotype but different underlying physiological states been experimentally identified? Ideally, we would like to find *E. coli* strains with different resource allocation strategies and corresponding biomass compositions, but with the same growth rate and growth yield. There are not many *E. coli* strains for which the biomass composition has been extensively characterized, but some partial information can be obtained from mutants which accumulate large amounts of storage metabolites, in particular glycogen. Under the assumption that the total biomass concentration, or biomass density, is approximately constant across different growth conditions [63, 64], a high glycogen mass fraction means that other mass fractions must be lower. This particularly concerns the mass fractions of protein and RNA, which account for more than 75% of total biomass.

As a first example, we combined two datasets obtained for the *E. coli* BW25113 strain [36, 37]. The first study quantified for each single-gene deletion mutant in the Keio collection [65], grown in minimal medium with glucose supplemented with amino acids, a large number of growth phenotypes, including growth rate and OD_600_ saturation level, a proxy for growth yield [36]. We combined this information with measurements of the amount of glycogen obtained in a second study. The authors of this study cultured mutants from the Keio collection in a medium with glucose and amino acids, and for each strain quantified the glycogen contents [37]. Whereas glycogen equals 0.026 mg glucose per mg protein in the wild-type strain, in some mutants the amount of glycogen increases up to ten-fold. We used this information to identify mutants having (i) a growth rate and a growth yield close to the average rate-yield phenotype of the entire strain collection, and (ii) a high amount of glycogen (at least 50% higher than the wild type).

Figure 7A singles out three mutant strains with a growth rate of 0.61 h^−1^ and a maximum OD_600_ level of 0.73, which are close to the average values and the values of the wild-type strain [36]. The strain with a deletion of *serA*, encoding an enzyme in the L-serine biosynthesis pathway, accumulates glycogen at a level of 0.23 mg glucose per mg protein, 9 times higher than in the wild-type strain [37]. Assuming that the biomass density remains approximately constant, the high concentration of glycogen must come at the expense of other biomass components, in particular protein and RNA. When trading the increase in glycogen mass against a proportional decrease in protein and RNA, the protein mass fraction of the ∆*serA* strain would decrease from 72% to 63% (*Methods*). This suggests that the average rate-yield phenotype can be achieved for quite different biomass compositions, resulting from different underlying resource allocation strategies.

**Figure 7.**
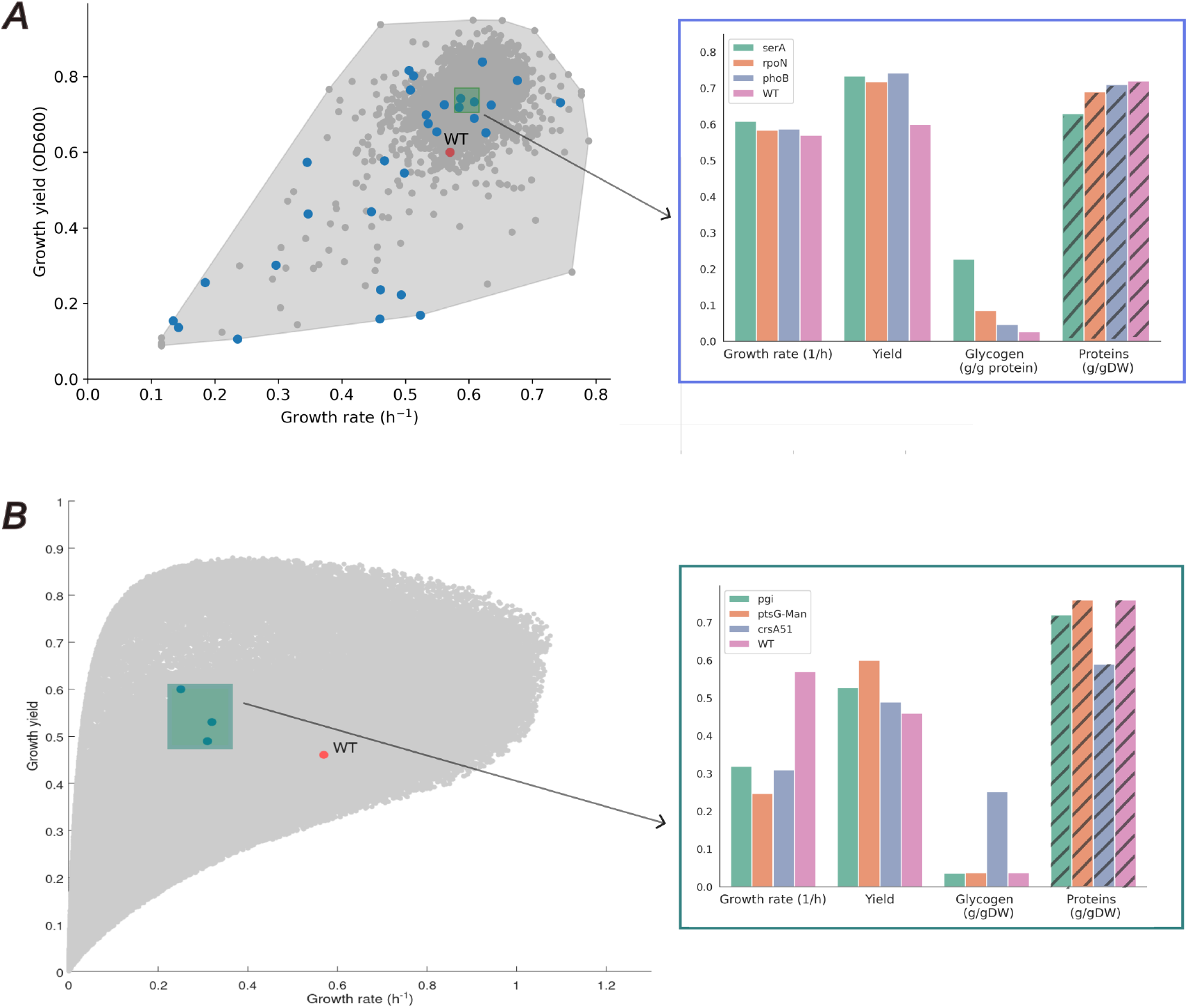
Comparison of biomass compositions of *E. coli* strains with high and low glycogen contents, but similar rate-yield phenotypes. (A) Three BW25113 deletion strains (∆*serA*, ∆*rpoN*, ∆*phoB* ) were found to have an average rate-yield phenotype (blue dots in green rectangle), but a higher glycogen contents than the wild-type strain, quantified in units mg glucose per mg protein (data from [36, 37]). The protein mass fractions (hatched) were inferred from the wild-type biomass composition and the measured glycogen contents under the assumption of constant biomass density (*Methods*). The rate-yield phenotype of the wild-type strain is shown for reference (red dot) against the background of all other mutant strains characterized in the study (grey dots). (B) The strains with deletions of *pgi* or *ptsG*-*malEFG* have a similar rate-yield phenotype as the *csrA51* mutant strain, but contrary to the latter, are not reported to have a glycogen excess [37, 38, 60, 66]. The protein mass fractions (hatched) were inferred from the data as in panel A. The rate-yield phenotype of the wild-type strain is shown for reference (red dot) against the background of predicted rate-yield phenotypes for this condition from Fig. 5B (grey dots).

A second example concerns the glycogen-overproducing *E. coli* mutant strain *csrA51* [38] (Figure 7B). The gene *csrA* encodes the global regulator CsrA and the mutant *csrA51* deletes the last 10 amino acids of the protein. The *csrA51* mutant has a large range of physiological effects, most importantly for our purpose, a strong increase in glycogen accumulation. In particular, *E. coli* cells grown in minimal medium with glucose were shown to multiply their glycogen concentration seven-fold, to a concentration of 0.25 g glucose per gDW. That is, 25% of the cellular biomass in this mutant consists of glycogen. By the same reasoning as above, one expects that the biomass fractions of the other cellular components, in particular protein and RNA, must decrease in proportion. More precisely, taking the protein and RNA mass fractions in the wild-type strain as reference, the protein mass fraction is estimated to decrease to 59% (Figure 7B).

The growth rate and growth yield of the *csrA51* mutant strain were measured as 0.31 h^−1^ and 0.49, respectively (Figure 7B, [38]). The growth rate is lower than that of the MG1655 wild-type strain by a factor of two, whereas the growth yield is similar. We searched within the database of rate-yield phenotypes used in Figure 5B for other *E. coli* strains that have approximately the same growth rate and growth yield as the *csrA51* mutant, but without excessive glycogen accumulation. We thus found, for example, the strain with a deletion of the glycolysis gene *pgi*, which has a growth rate and growth yield of 0.32 h^−1^ and 0.53, respectively (Figure 7B, [60]). Since this mutant does not produce glycogen in excess [37], the biomass composition of the strain is likely to resemble that of the wild-type strain, that is, high protein and low glycogen mass fractions. Again, this suggests that different biomass compositions, and different underlying resource allocation strategies, may give rise to the same rate-yield phenotype.

### Experimental evidence for alternative physiological states in microalgae

Whereas the total protein concentration of *E. coli* generally does not vary much across standard laboratory conditions, greater variability in biomass composition is observed in microalgae. The protein mass fraction typically ranges from 30 to 65%, while carbohydrate and lipid fractions range from 5 to 45% and 15 to 65%, respectively [67]. The latter two fractions include both structural and storage components. In particular, storage metabolites are typically accumulated under nutrient starvation.

The question can be asked whether the consequences of suboptimality outlined above also apply to organisms like microalgae, which regulate their biomass composition quite differently. A recent study [39] characterized a hundred clones of the haptophyte *Tisochrysis lutea*, obtained by isolating clones from 15 polyclonal strains. Nitrogen-limited batch cultures were conducted, during which growth rates (measured in exponential phase) and final (carbon) biomass were quantified; the latter is hereafter used as yield. In addition, protein and total fatty acids contents were measured at the end of the culture. The latter may serve as a proxy for carbon storage, although this measure likely also includes a fraction of functional (non-storage) lipids. Other forms of carbon storage, such as carbohydrates, are also possible in microalgae but were not measured in this study.

Figure 8 shows the relationship between growth yield and growth rate for the different *T. lutea* clones. The convex envelope of the data can be used as an approximation of a Pareto front to delineate a set of suboptimal clones, defined as those located within the interior, away from the hypothetical Pareto front. Interestingly, among these suboptimal clones, individuals with nearly identical yield and growth rate can display markedly different biomass compositions. We selected clones with a growth rate around 0.5 d^−1^ and a growth yield around 0.4 gC L^−1^, the population means for these two quantities. For example, clone Tiso-3-Cl1 exhibits approximately 50% higher protein content but roughly twofold lower total fatty acids content than a clone of similar performance (Tiso-12-Cl6). Overall, for clones with similar rate-yield phenotypes, the protein and fatty acids contents are negatively correlated, as expected when the total biomass concentration is approximately constant (Figure 8). The suboptimal clones reflect distinct resource allocation strategies, despite sharing comparable growth rates and yields, thus extending the conclusions for *E. coli* to protists.

**Figure 8.**
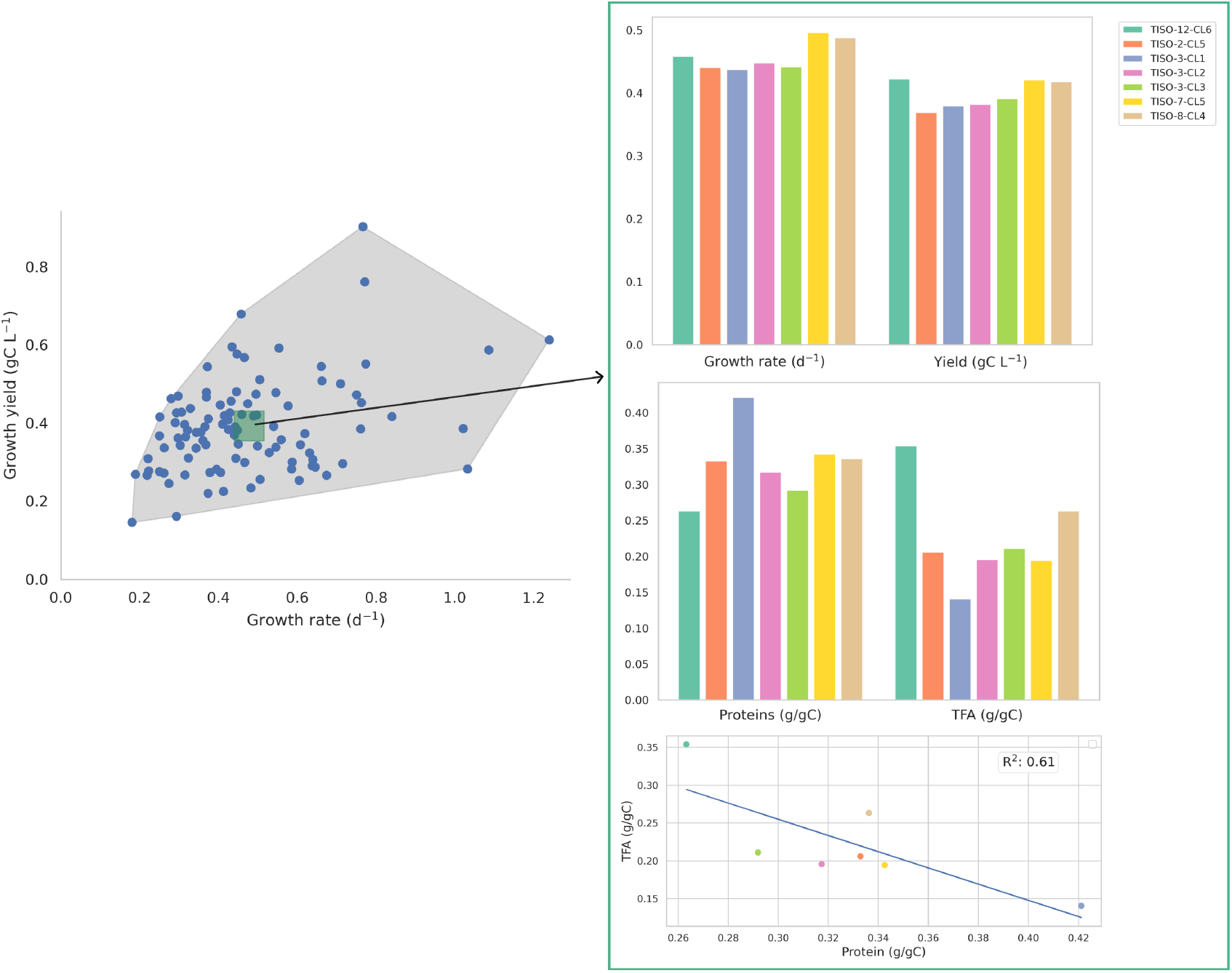
Comparison of the biomass composition of *T. lutea* strains with variable total protein and fatty acids contents, but similar rate-yield phenotypes. Measured growth yields and growth rates for different *T. lutea* clones [39]. The biomass composition of clones with similar suboptimal rate-yield phenotypes is compared, showing highly variable and negatively correlated total protein and fatty acids contents (R^2^ = 0.61).

## Discussion

The assumption that organisms have evolved to optimize a certain phenotypic characteristic or combination of characteristics is common in the analysis of the behavior of microorganisms and other biological systems. The optimality assumption has led to good results when studying microbial growth, but there are also counterexamples where it does not seem to hold. Among the reasons for observed deviations from predicted optimality, which have been extensively discussed in the literature relating optimality and natural selection [1, 2, 3, 4, 5, 6, 7], we mentioned here an inappropriate choice of supposedly optimized phenotypic characteristics and a mismatch between laboratory conditions and environmental niches in which microorganisms have evolved. While recent work on inverse optimality [68], where the objective functions are also inferred from observations of the biological system, goes some way in addressing the above issues, it may not completely counter all arguments against optimality.

In the present paper, we have deliberately not taken position in this debate. Instead, we started from the reasonable assumption that in many practical situations of interest, microbial growth cannot be assumed Pareto optimal with respect to commonly considered characteristics like growth rate and growth yield. This has motivated the analysis of resource allocation and microbial growth under the assumption of Pareto suboptimality. Whereas a general theory predicts the existence of a Pareto frontier from properties of the mapping from cellular resource allocation to phenotypes [8, 9, 40], here summarized as Theorem 1, much less is known about the relation between resource allocation and Pareto suboptimal phenotypes.

Suboptimal resource allocation has usually been treated from the perspective of trade-offs, where suboptimal performance with respect to one phenotypic characteristic is accompanied by superior performance with respect to another phenotypic characteristic [11]. For example, the constitution of a ribosome reserve may be detrimental to growth rate, but speeds up cellular response when the growth conditions improve [26, 69]. Given the linear relation observed between growth rate before starvation and death rate during starvation, a suboptimal growth rate when nutrients are abundant leads to longer survival when nutrients are depleted [70]. The adaptation to a new environment may not be optimal in some conditions, but it avoids regulatory complexity (and its associated costs) that would come with the required finetuning of gene expression [52]. These examples illustrate the point of Fig. 3 that modifying the set of phenotypic characteristics under consideration may bring a suboptimal phenotype closer to the Pareto frontier.

In comparison, we have taken a different perspective here, focusing on the consequences of the location of phenotypes away from the Pareto frontier. Under the same general assumptions as the theoretical results mentioned above, we demonstrated that there exist multiple resource allocation strategies mapping to the same Pareto suboptimal phenotype (Proposition 2), contrary to Pareto optimal phenotypes which arise from a unique strategy. Each of the suboptimal strategies is suboptimal in a different way, incurring different so-called growth costs [54, 71]. For example, ribosomes can be expressed in excess of what is needed for protein synthesis, thus reducing the resources available for enzymes in central carbon and energy metabolism. Alternatively, producing ATP in excess by investing a larger fraction of resources in respiratory enzymes may come at the cost of lower ribosome expression, thus leaving ATP underutilized for protein synthesis and increasing ATP spillage. While exhibiting the same phenotype, these suboptimal strategies lead to physiological states, and corresponding biomass compositions, that may be quite different, especially as the distance to the Pareto frontier increases.

Inspired by the mathematical results on Pareto suboptimality, we analyzed microbial growth using a coarse-grained model of coupled carbon and energy fluxes, calibrated for *E. coli* [29]. Despite its limitations, including the restriction of resource allocation constraints to the distribution of protein synthesis capacity over different protein classes and the absence of regulation of enzyme activity, the model was shown to agree quite well with a database of rate-yield phenotypes reported for *E. coli* strains. We extended the model to take into account the synthesis of storage metabolites, in particular glycogen. Analysis of the model predictions confirmed, as expected from theory, that a given suboptimal rate-yield phenotype can be achieved by means of a whole range of resource allocation strategies. Interestingly, some of the biomass compositions corresponding to these strategies show large variations in the total concentration of storage metabolites, at the expense of the total protein concentration. This is reminiscent of experimental observations that intracellular carbon storage by microorganisms is a pathway to biomass accumulation alternative to cellular replication [72].

A rigorous test of the predictions of the model would require integrated datasets of physiology, proteomics, and metabolomics, which are not available. Nevertheless, by combining datasets from the literature with suitable assumptions, we were able to make some interesting observations. First, we identified several *E. coli* strains with similar, suboptimal rate-yield phenotypes but very different glycogen concentrations. Under the assumption that the biomass density remains approximately constant across different strains, well-supported by the available experimental data [63, 64], we concluded that the five to ten-fold higher higher glycogen mass fraction must be compensated for by significant decreases in the mass fraction of other biomass components, notably protein.

Second, we complemented this observation with recently published measurements of the biomass composition of different clones of the microalga *T. lutea*. In this case, we also found that the same suboptimal rate-yield phenotype could be obtained with very different biomass compositions. Moreover, the protein and fatty acids mass fractions were negatively correlated, thus indicating that an increase of the latter requires a decrease of the former, as expected from the assumption of constant biomass density. The data thus confirm that the same suboptimal phenotype can be achieved for a range of different biomass compositions associated with different resource allocation strategies, as predicted by the theory (Proposition 2).

The argument that a given Pareto suboptimal phenotype can be achieved for a range of resource allocation strategies, corresponding to different biomass compositions, raises another question. It suggests that suboptimal strategies enable a diversity of responses to changes in the environment. For example, while microorganisms display the same growth rate and growth yield, the investment of underutilized resources in a variety of cellular functions could lead them to respond differently when faced with an external stress, such as a change in nutrient availability. This association of suboptimality with diversity brings us back to the discussion of trade-offs above, but from a slightly different angle. Instead of trade-offs between specific phenotypic characteristics, microorganisms may follow a more general trade-off strategy, exploiting the fact that suboptimal performance on any one characteristic releases resources that can be used to sustain a diversity of performances on any other characteristic.

## Methods

### Precise statement and proof of mathematical results

We formulate and prove mathematical results for a generalization of the multiobjective optimization problem of Eq. 1. These results imply Theorem 1 and Proposition 2.

### Basic definitions

#### Definition 3

A multiobjective optimization problem has the form

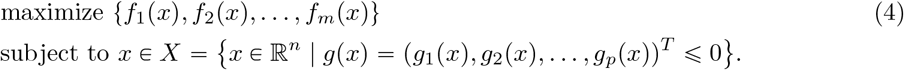

Variables *x*_*i*_ are decision variables, functions *f*_*j*_ objective functions, functions *g*_*k*_ constraints, and set *X* is the feasible set. The set *P = f (X)* of all possible objective values is called the objective set.

Note that this definition generalizes the multiobjective optimization problem of Eq. 1 to *p ≥* 2 objectives.

#### Definition 4

A function *h* : *X* → ℝ is convex if for all *x, y* ∈ *X* and all 0 ≼ *β* ≼ 1

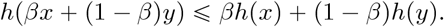

A set *S* ⊂ ℝ^*n*^ is convex if for any *x, y* ∈ *S* we have

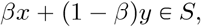

for all 0 ≼ *β* ≼ 1.

A function *f* : *X* → ℝ is strongly convex if there exists *α* > 0 such that

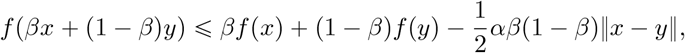

for all 0 ≼ *β* ≼ 1.

#### Definition 5

A multiobjective optimization problem is convex if all the objective functions and the feasible objective region *X* are convex.

The solution *v* = (*f*_1_, *f*_2_, …, *f*_*m*_) ∈ *P* dominates solution *u* = (*f*_1_′, *f*_2_′, …, *f*_*m*_′ ) ∈ *P*, written as *v* > *u*, if

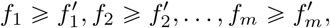

and at least one of the inequalities is strict.

#### Definition 6

(**Pareto optimality**). An objective vector *v*^*^ ∈ *P* is Pareto optimal if there does not exist another objective vector *v* ∈ *P* such that *v* dominates *v*^*^. The Pareto frontier *P* ^*^ is the set of Pareto optimal objective vectors *u* ∈ *P* . In other words,

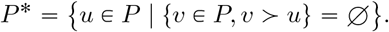

A decision vector *x*^*^ ∈ *X* is Pareto optimal if there does not exist another decision vector *x* ∈ *X*such that *f* (*x*) > *f* (*x∈*). The Pareto set *L* ⊂ *X* is the set of decision vectors *x* such that the value of the function *f* (*x*) ∈ *P* ^*^ belongs to Pareto frontier. In other words, *f* (*L*) = *P* ^*^.

### Proof of Theorem 1

A special case of Theorem 1 was proved by Shoval *et al*. [8] and then proved in full generality by Hamada *et al*. [40]. The proof in this section follows the latter exposition. For clarity of exposition, we repeat the theorem first.

#### Theorem 1

Provided that

Assumption 1 the objective functions *f*_1_, *f*_2_ and the constraint functions *g*_*i*_ are continuously differentiable,

Assumption 2 each function *f*_1_, *f*_2_ has a unique maximum in the interior of the feasible set *X*, obtained for strategies *m*_1_, *m*_2_,

Assumption 3 the maxima are different, *i*.*e*., *m*_1_ ≠ *m*_2_,

Assumption 4 the functions *f*_1_, *f*_2_ and the set *X* are strongly convex, it holds that

1. there exists a Pareto frontier *P* ^*^,
2. the Pareto frontier *P* ^*^ is a curve that joins the maximum of *f*_1_ to the maximum of *f*_2_,
3. the multiobjective optimization problem of Eq. 1 can be formulated as a collection of single objective optimization problems

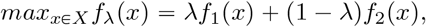

where 0 *≼ λ ≼* 1 indicates the relative preference for *f*_1_ and *f*_2_. For well-behaved problems that satisfy the above assumptions, the Pareto frontier *P* ^*^ will be exactly the set of optima for the above problem as the value of *λ* changes between 0 and 1.

Conclusion 1 of the theorem follows from compactness of the set *X* and continuity of *f* . Indeed, the image *P f X* of a compact set *X* by a continuous function *f* is compact and therefore bounded. In order to prove conclusions 2 and 3, we introduce the concept of weakly simplicial optimization problem for functions *f* : ℝ^*n*^ → ℝ^+1^ [40].

#### Definition 7

A standard m-simplex is a set

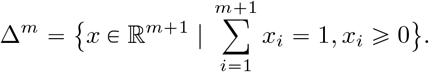

We define the index set *M* = {1,…,*m* + 1 } and denote a face of ∆^*m*^ for *I* ⊆ *M* by

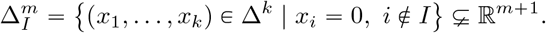

#### Definition 8

Let *X* be a subset of ℝ^*n*^ and *f* = (*f*_1_, …, *f*_*m*+1_) be a mapping *f* : *X* → ℝ^*m*+1^. For *I = { i*_1_, …, *i*_*k*_ *} M* such that *i*_1_ *i*_*k*_, we put *f*_*I*_ *= (f*_*i*1_, …, *f*_*ik*_ ).

The problem of minimizing *f* is *C*^*r*^-weakly simplicial if there exists a *C*^*r*^-mapping *ϕ* : ∆^*m*^ *P* satisfying

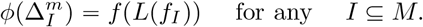

#### Theorem 9

(Theorem 1.1 in [40]). Let *f* : ℝ^*n*^ ℝ^*m*+1^ be a strongly convex *C*^*r*^-mapping with ( 2 ≼ ≼ ∞) . Then the multiobjective optimization problem of minimizing *f* is *C*^*r* -1^-weakly simplicial (Figure 9).

**Figure 9.**
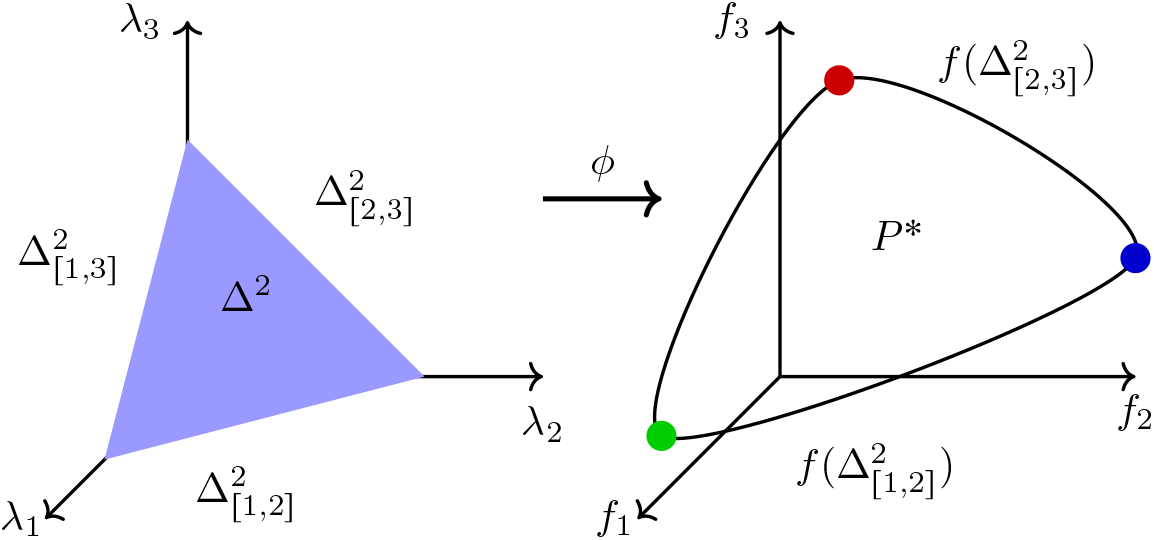
Example of a weakly simplicial multiobjective optimization problem, illustrating Theorem 9 [40]. Consider the weakly simplicial multiobjective optimization problem maxt*f*_1_, *f*_2_, *f*_3_u. On the left is a 2 simplex ∆ of parameters *λ* turning the multiobjective optimization problem to a single-objective optimization problem *f*_*x*_ = *λ*_1_*f*_1_ + *λ*_2_*f*_2_ + *λ*_3_*f*_3_. A weakly simplicial problem has a smooth map *ϕ* which not only maps ∆ to the Pareto frontier *P*, but every sub-simplex on the left to a corresponding subset of the Pareto frontier on the right. The maxima of functions *f*_1_ (green), *f*_2_ (blue) and *f*_3_ (red) are indicated.

We apply the theorem to the multiobjective optimization problem of Eq. 1. We note that maximization of *f* = (*f*_1_, *f*_2_) is equivalent to minimization of (−*f*_1_, −*f*_2_), and hence Theorem 9 applies to the problem. Assumptions 2 and 3 of Theorem 1 ensure that the maxima of *f*_1_ and *f*_2_ are in generic positions within the set *X*. Then, in view of Assumption 4, Theorem 9 implies that the Pareto frontier *P* ^*^ ⊂ *P* is a smooth image of an interval (1-simplex) connecting the maximum of *f*_1_ and the maximum of *f*_2_. This proves conclusion 2 of Theorem 1.

Furthermore, Proposition 2.5 in [40] shows that under the assumptions of Theorem 9, the weighted-sum scalarization *w* = (*w*_1_, …, *w*_*m*+1_) ∈ ∆^*m*^, such that _*i*_ *w*_*i*_ = 1, gives an instance of the required homeomorphism *ϕ*. In other words, the mapping *ϕ* : ∆^*m*^ → *P* defined by

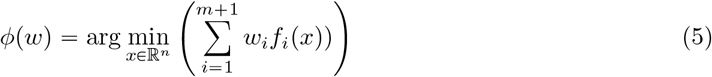

is a diffeomorphism. Conclusion 3 directly follows from Eq. 5.

**Proof of Proposition 2**

#### Proposition 2

Retain Assumptions 1-4 from Theorem 1 and make the further assumption that the Pareto set lies in the interior of *X*. Then, for any feasible phenotype dominated by the Pareto frontier *P* ^*^, *p* ∈ *P* \*P* ^*^, and lying in the neighborhood of *P* ^*^, there exist feasible *x, x*^′^ ∈ *X, x* ≠ *x*^′^, such that (*f*_1_(*x*), *f*_2_(*x*)) = (*f*_1_(*x*^′^), *f*_2_(*x*^′^)) = *p*.

The discussion in the main text provides the intuition for this result in case we have two decision variables *x* (Figure 4). However, for more general multiobjective optimization problems, with more than two decision variables and possibly also more than two objective functions (Definition 4), there generally exists an infinite number of equivalent strategies *x* and *x*^′^. This follows from standard differential geometry results which we briefly summarize below.

We consider a general differentiable *C*^1^ map *f* : *M N* where *M, N* are differentiable *C*^1^ manifolds [73].

#### Definition 10

Let *f* : *M* → *N* be a *C*^1^ map. We call *x* ∈ *M* a regular point if *T*_*x*_*f* : *M*_*x*_ → *N*_*f*p*x*q_ is a surjective linear map from tangent space of *M* at *x* to tangent space of *N* at *f* (*x*). Otherwise *x* is a critical point and *f* (*x*) is a critical value. If *y* ∈ *N* is not a critical value, then *f* ^−1^(*y*) is a regular value, even if *y* R *f* (*M* ).

Note that a Pareto frontier *P* ^*^ consists of critical values of *f* since the gradients of the objective functions ∇*f*_*i*_ *x* are linearly dependent at *x ∈ L*. However, the Morse-Sard theorem shows that the set of regular values for sufficiently smooth functions is dense in *N* and have full measure and therefore almost all values away from Pareto front are regular values.

#### Theorem 11

(Morse-Sard Theorem, [73]). Let *M, N* be manifolds of dimension *m, n* respectively and let *f* : *M → N* be a *C*^*r*^ function with *r >* max {0, *m n}*. Then the set of critical values has zero measure in *N* . Furthermore, the set of regular values is residual and therefore dense in *N* .

In addition, the inverse images of the regular values have predictable structure.

#### Theorem 12

(Theorems 3.2 and 3.3 in [73]). Let *f* : *M → N* be a *C*^*r*^, *r ≽* 1 map. If *y* ∈ *f(M)* is a regular value, then *f* ^-1^ *y* is a *C*^*r*^-submanifold of *M* with the same codimension in *M* as the dimension of *N* .

From the above follows a corollary that directly implies Proposition 2 in the main text.

#### Corollary 13

Consider the multiobjective optimization problem defined by *f* : ℝ^*n*^ → ℝ^*m*^ with *n* decision variables and *m* objective functions with *k* ≼ *m* (Eq. 1). If *p* is a regular value of *f*, then the set of equivalent decision vectors *x* ∈ *X* supporting phenotype *p* ∈ *P* is a sub-manifold of *X* of dimension *n−m*.

When applied to the bacterial growth example, where *n* 3 and *m* 2, the set of equivalent resource allocation strategies must be a sub-manifold of *X* of dimension 1. Figure 6B shows this is indeed the case.

### Model of bacterial growth

The present model is an extension of a previous resource allocation model describing the coupling of carbon and energy fluxes in microbial growth, in particular *E. coli* [29]. The equations are summarized in *Text S1*. The main change with respect to the previous model is the addition of a macroreaction for the synthesis of glycogen, a storage metabolite, from precursor metabolites, which consumes ATP in the process. The reaction is assumed to follow irreversible Michaelis-Menten kinetics, like the other reactions in the model, and is catalyzed by enzymes from central carbon metabolism. The fraction of carbon and energy source converted into glycogen is usually low, but when the supply of carbon source exceeds the demand for macromolecular synthesis, this fraction starts to increase. This involves complex regulatory mechanisms of the affinity and rate of a key enzyme in the glycogen synthesis pathway in *E. coli* [56, 74, 75], which we approximated here by choosing a high value for the half-saturation constant of the reaction. More generally, model calibration followed the same procedure as for the original model (*Text S1*, [29]). The procedure does not require computational parameter fitting, since all parameters are unambiguously fixed by data from the literature and suitable hypotheses motivated by experimental results. The model file and the Matlab R2025 code to calibrate and run the model are available from https://gitlab.inria.fr/baldazzi/glycogen-model.

### Experimental data on *E. coli* and *T. lutea*

The following data from the literature were used in this study: measurements of growth rate (min^−1^) and maximum OD_600_ (growth yield) of *E. coli* strains from the Keio collection cultured in minimal medium with glucose supplemented with amino acids [36, 65]; measurements of glycogen concentrations 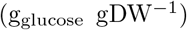of *E. coli* strains from the Keio collection cultured in minimal medium with glucose supplemented with amino acids [37, 65]; measurements of growth rate (h^−1^),growth yield 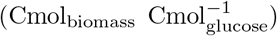, and glycogen concentration (g_glucose_ gDW^−1^) of *E. coli csrA51* strain cultured in minimal medium with glucose [38]; a compilation of measurements of growth rate and growth yield of *E. coli* strains cultured in minimal medium with glucose [29]; measurements of growth rate (d^−1^), total content of carbon biomass (gC L^−1^, growth yield), total protein concentration (g gC^−1^), and total fatty acids concentration (g gC^−1^) of different clones of the algae *T. lutea* [39]. The experimental data are available as *File S1* and the code for analyzing the data is available from https://gitlab.inria.fr/baldazzi/glycogen-model.

In order to estimate the change in biomass composition to compensate for the increase in glycogen contents in the *E. coli* strains, under the constraint of constant total biomass density, we used the formula *f*_*p*_ + *f*_*u*_ + *f*_*c*_ + *f*_*g*_ = 1, where *f*_*p*_ is the protein mass fraction, *f*_*u*_ the mass fraction of other macromolecules, *f*_*c*_ the mass fraction of metabolites in central carbon metabolism, and *f*_*g*_ the glycogen mass fraction. Measurements of glycogen contents are usually reported in units 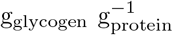, that is, as a fraction *f*_*g,p*_ of total protein [37], so that *f*_*g*_ = *f*_*g,p*_ *f*_*p*_ and the mass conservation equation can be written as (1 + *f*_*g,p*_) *f*_*p*_ + *f*_*u*_ + *f*_*c*_ = 1. Consider a mutant strain for which the value of *f*_*g,p*_ is higher than that of the wild-type strain. When the values of *f*_*p*_, *f*_*u*_, and *f*_*c*_ are not known, we make the assumption that they are proportionally decreased by a factor *λ* with respect to the known values of the wild-type strain to maintain constant biomass density. That is, we write 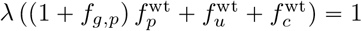, where the subscript denotes the known values of the mass fractions for the wild-type strain. After solving for *λ*, we can compute the estimated mass fractions for the different biomass components of the mutant strain as 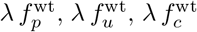, and 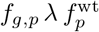. If the glycogen contents are reported in units g_glycogen_ gDW^−1^, that is, as a fraction *f*_*g*_ of total biomass [38], the computation of the proportionality factor needs to be slightly modified. Analogously to the reasoning above, we then need to solve *λ* from 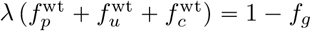

## Supplementary information

**Text S1** Coarse-grained resource allocation model of microbial growth.

**File S1** Compilation of experimental data used in this study.

## Acknowledgments

This work has been supported by the ANR (CTRL-AB project, ANR-20-CE45-0014), PEPR B-BEST “Biomasses, biotechnologies et technologies durables pour la chimie et les carburants” (MuSiHC project, ANR-24-PEBB-0006), PEPR AgroEcoNum (MISTIC project, ANR-22-PEAE-0011), and by Centre Inria de l’Université Grenoble Alpes (funding for visits of Tomas Gedeon). The authors would like to thank Mathieu Besançon for discussions.

## Notes

### Competing Interest Statement

The authors have declared no competing interest.

