## Supplemental Text 1 for "Pareto Suboptimal Resource Allocation and Microbial Growth"

### Text S1: Coarse-grained resource allocation model of microbial growth*

Valentina Baldazzi[1,2,+], Francis Mairet[3], Tomas Gedeon[4,*], and Hidde de Jong[5,6,*,+]

[1]Université Côte d'Azur, Inria, INRAE, CNRS, Sophia Antipolis, France
[2]Université Côte d'Azur, INRAE, CNRS, Institut Sophia-Agrobiotech, Sophia Antipolis, France
[3]IFREMER, Phytox, Nantes, France
[4]Montana State University, Bozeman, United States, (0000-0001-5555-6741)
[5]Université Grenoble Alpes, Inria, Grenoble, France
[6]Université Grenoble Alpes, CNRS, LIPhy, Grenoble, France
[*]These authors contributed equally

September 7, 2026

#### Model description and model equations

The present model is an extension of a previous resource allocation model describing the coupled carbon and energy fluxes in microbial growth [1]. Briefly, the latter model is based on the partitioning of the biomass into proteins, other macromolecules like DNA and RNA, and central metabolites used for synthesis of the macromolecules. The proteome is divided into five categories responsible for the uptake and metabolism of external

*Suppementary information for "Pareto Suboptimal Resource Allocation and Microbial Growth"

substrates, protein synthesis, energy metabolism (via respiration or fermentation), and housekeeping functions, including the synthesis of other macromolecules. Allocation of resources to these different categories of proteins is governed by resource allocation parameters $x_c, x_r, x_{er}, x_{ef}, x_u$, respectively, which are positive and sum up to 1. Energy cofactors, ADP and ATP, are continuously recycled through the energy metabolism and the biosynthesis pathways (synthesis of proteins and other macromolecules). Their synthesis from central metabolites is not modelled explicitly; their total intracellular concentration is considered constant.

The previous model did not explicitly distinguish the presence of storage metabolites, like glycogen, and included them within the central metabolites. We modified the model to make this component explicit and defined the total cellular biomass $B$ [gDW] as

$$B = \beta\,(R + M_c + M_{er} + M_{ef} + M_u + C + G + U), \tag{S1}$$

where $1/\beta$ is the total biomass carbon content [Cmmol gDW$^{-1}$], $G$ the glycogen mass, $C$ the mass of the central carbon metabolites, $U$ the mass of other macromolecules, and $M_c, M_u, R, M_{er}, M_{ef}$ the mass of proteins associated with the different categories mentioned above. Note that, like in the original model, ATP and ADP are not included in the biomass, due to their negligible quantity.

Assuming that the volume of the growing microbial population is proportional to the biomass [2], we transform the above quantities into concentrations by dividing with the total biomass $B$: $m_u = M_u/B$, $m_c = M_c/B$, $m_{er} = M_{er}/B$, $m_{ef} = M_{ef}/B$, $r = R/B$, $c = C/B$, $g = G/B$, $u = U/B$. Accordingly, the concentration variables have units Cmmol gDW$^{-1}$ and the total biomass concentration, or biomass density, is given by $1/\beta$.

Glycogen is synthesized from central carbon metabolites and ATP, when an excess of carbon source is available [3, 4]. The corresponding reaction with rate $v_g$ [Cmmol gDW$^{-1}$ h$^{-1}$] is essentially irreversible under the growth conditions considered (batch growth in minimal medium with glucose) and is assumed to be controlled by the enzymatic pool $m_c$:

$$v_g = m_c\, k_g\, \frac{c}{c + K_g}\, \frac{a^*}{a^* + K_{ag}}, \tag{S2}$$

where $k_g$ is the maximum rate and $K_g$ and $K_{ag}$ are half-saturation constants for $c$ and $a^*$ substrates, respectively.

Like in the previous model [1], the dynamics of concentration variables can be described by a system of ordinary differential equations, which now includes an equation for glycogen:

$$\frac{dc}{dt} = v_{mc} - v_{mer} - \rho_{mef}\, v_{mef} - \rho_{ur}\, v_r - \rho_{ur}\, v_{mu} - v_g - (\mu + \gamma)\, c, \quad \text{(S3)}$$

$$\frac{du}{dt} = v_{mu} - (\mu + \gamma)\, u, \quad \text{(S4)}$$

$$\frac{dg}{dt} = v_g - (\mu + \gamma)\, g, \quad \text{(S5)}$$

$$\frac{dm_u}{dt} = x_u\, v_r - (\mu + \gamma)\, m_u, \quad \text{(S6)}$$

$$\frac{dr}{dt} = x_r\, v_r - (\mu + \gamma)\, r, \quad \text{(S7)}$$

$$\frac{dm_c}{dt} = x_c\, v_r - (\mu + \gamma)\, m_c, \quad \text{(S8)}$$

$$\frac{dm_{er}}{dt} = x_{er}\, v_r - (\mu + \gamma)\, m_{er}, \quad \text{(S9)}$$

$$\frac{dm_{ef}}{dt} = x_{ef}\, v_r - (\mu + \gamma)\, m_{ef}, \quad \text{(S10)}$$

(S11)

where the reaction rates $v_{mc}, v_{mer}, v_{mef}, v_{mu}, v_r$ [Cmmol gDW$^{-1}$ h$^{-1}$] and $v_d$ [mmol gDW$^{-1}$ h$^{-1}$] are defined as follows (for $v_g$, see Eq. (S2) above):

$$v_{mc} = m_c\, e_m, \quad \text{(S12)}$$

$$v_{mer} = m_{er}\, k_{mer} \frac{c}{c + K_{mer}} \frac{a_0 - a^*}{a_0 - a^* + K_{amer}}, \quad \text{(S13)}$$

$$v_{mef} = m_{ef}\, k_{mef} \frac{c}{c + K_{mef}} \frac{a_0 - a^*}{a_0 - a^* + K_{amef}}, \quad \text{(S14)}$$

$$v_{mu} = m_u\, k_{mu} \frac{c}{c + K_{mu}} \frac{a^*}{a^* + K_{amu}}, \quad \text{(S15)}$$

$$v_r = r\, k_r \frac{c}{c + K_r} \frac{a^*}{a^* + K_{ar}}, \quad \text{(S16)}$$

$$v_d = k_a\, a^*. \quad \text{(S17)}$$

In addition, the energy and mass flows are coupled via the following balance equation, now accounting for the ATP cost of glycogen synthesis:

$$\frac{da^*}{dt} = n_{mer}\, v_{mer} + n_{mef}\, v_{mef} - n_r\, v_r - n_{mu}\, v_{mu} - n_g v_g - v_d, \quad \text{(S18)}$$

where $a^*$ [mmol gDW$^{-1}$] is the ATP concentration, and $a_0 - a^*$ the ADP concentration, with $a_0$ the total (constant) concentration of ATP and ADP.

Note that $\mu$ [h$^{-1}$] denotes the growth rate of the population and $\gamma$ [h$^{-1}$] the biomass degradation rate. Like in the original publication [1], the growth rate is expressed as a function of the reaction rates as follows:

$$\mu = \frac{1}{B}\frac{dB}{dt} = \beta\,(v_{mc} - v_{mer} - \rho_{mef}\,v_{mef} - (\rho_{ur} - 1)\,(v_{mu} + v_r)) - \gamma. \quad \text{(S19)}$$

The nondimensional growth yield $Y$ is defined as the ratio of the net biomass synthesis rate ($\mu/\beta$) and the carbon uptake rate $v_{mc}$, which leads to the following expression:

$$Y = \frac{1}{\beta}\frac{\mu}{v_{mc}} = \frac{v_{mc} - v_{mer} - \rho_{mef}\,v_{mef} - (\rho_{ru} - 1)\,(v_r + v_{mu}) - \gamma/\beta}{v_{mc}}. \quad \text{(S20)}$$

The definition of all variables and parameters is summarized in Tables S1.1 and S1.2.

#### Model calibration

Model calibration follows the same steps as in the original publication [1] and was performed using published reference datasets with measurements of growth rates and fluxes [5, 6, 7], protein concentrations [8], and metabolite concentrations [6, 9, 10] for the *E.coli* BW25113 strain under steady-state, exponential (batch) growth in minimal medium with glucose. Accordingly, most of the estimated concentration values and kinetic parameters coincide with those in [1] and are summarized in Tables S1.3 and S1.4. Below, we focus the discussion on the calibration of the glycogen synthesis reaction.

The calibration procedure starts by using the total biomass density and measured biomass fractions of proteins and metabolites to derive total protein and metabolite concentrations. Storage metabolites can be derived from a literature study [3], which performed a systematic quantification of glycogen content in BW25113 wild-type and mutant strains. For the wild-type strain, the average glycogen content was equivalent to 147 nmol glucose/mg protein, which equals 0.026 g glc/g protein. Based on the data from [2] and observed growth rates of the BW25113 strain in glucose minimal medium, we previously estimated the protein dry mass fraction to be 0.72 [1]. Assuming a glycogen structure $(C_6H_{10}O_5)_n$ with $n = 10$ on average [4], we can estimate the glycogen concentration as:

$$\hat{g} = (0.026/0.72) \cdot (60 \cdot 1000/1640) = 1.32 \text{ Cmmol gDW}^{-1}, \quad \text{(S21)}$$

| Model | Description | Unit |
|---|---|---|
| | **Macromolecule concentrations** | |
| $r$ | ribosomes | Cmmol gDW$^{-1}$ |
| $m_c$ | enzymes in central carbon metabolism | Cmmol gDW$^{-1}$ |
| $m_{er}$ | enzymes in energy metabolism (respiration) | Cmmol gDW$^{-1}$ |
| $m_{ef}$ | enzymes in energy metabolism (fermentation) | Cmmol gDW$^{-1}$ |
| $m_u$ | other proteins | Cmmol gDW$^{-1}$ |
| $u$ | other macromolecules | Cmmol gDW$^{-1}$ |
| | **Metabolite concentrations** | |
| $c$ | central carbon metabolites | Cmmol gDW$^{-1}$ |
| $g$ | storage metabolites (glycogen) | Cmmol gDW$^{-1}$ |
| $a^*$ | ATP | mmol gDW$^{-1}$ |
| | **Reaction rates** | |
| $v_{mc}$ | carbon uptake and central metabolism | Cmmol gDW$^{-1}$ h$^{-1}$ |
| $v_g$ | glycogen synthesis | Cmmol gDW$^{-1}$ h$^{-1}$ |
| $v_{mer}$ | energy metabolism (respiration) | Cmmol gDW$^{-1}$ h$^{-1}$ |
| $v_{mef}$ | energy metabolism (fermentation) | Cmmol gDW$^{-1}$ h$^{-1}$ |
| $v_r$ | protein synthesis | Cmmol gDW$^{-1}$ h$^{-1}$ |
| $v_{mu}$ | synthesis of other macromolecules | Cmmol gDW$^{-1}$ h$^{-1}$ |
| $v_d$ | energy dissipation | mmol gDW$^{-1}$ h$^{-1}$ |
| | **Other rates and yield** | |
| $\mu$ | growth rate | h$^{-1}$ |
| $\gamma$ | degradation rate | h$^{-1}$ |
| $Y$ | growth yield | - |

**Table S1.1: Model variables and rates.** The units Cmmol and gDW refer to mmol carbon and gram dry weight, respectively.

| Model | Description | Unit |
|---|---|---|
| | **Resource allocation parameters** | |
| $x_r$ | fraction of ribosomal proteins | - |
| $x_c$ | fraction of enzymes in central carbon metabolism | - |
| $x_{er}$ | fraction of enzymes in respiration | - |
| $x_{ef}$ | fraction of enzymes in fermentation | - |
| $x_u$ | fraction of other proteins | - |
| | **ATP factors** | |
| $n_{mer}$ | ATP yield from respiration | mmol $Cmmol^{-1}$ |
| $n_{mef}$ | ATP yield from fermentation | mmol $Cmmol^{-1}$ |
| $n_r$ | ATP cost of protein synthesis | mmol $Cmmol^{-1}$ |
| $n_{mu}$ | ATP cost of synthesis of other macromolecules | mmol $Cmmol^{-1}$ |
| $n_g$ | ATP cost of glycogen synthesis | mmol $Cmmol^{-1}$ |
| | **Correction factors** | |
| $\rho_{mef}$ | correction for $CO_2$ loss during fermentation | - |
| $\rho_{ru}$ | correction for $CO_2$ loss during macromolecular biosynthesis | - |
| | **Kinetic parameters** | |
| $k_{mef}$, $k_{mer}$, $k_r$, $k_{mu}$, $k_g$, $e_s$ | maximum rate of (enzymatic) reaction | $h^{-1}$ |
| $k_a$ | ATP dissipation rate | $h^{-1}$ |
| $K_{mef}$, $K_{mer}$, $K_r$, $K_{mu}$, $K_g$ | half-saturation for central metabolites | Cmmol $gDW^{-1}$ |
| $K_{amef}$, $K_{amer}$, $K_{ar}$, $K_{amu}$, | half-saturation for ATP | mmol $gDW^{-1}$ |
| | **Other constants** | |
| $1/\beta$ | total biomass concentration | Cmmol $gDW^{-1}$ |

**Table S1.2: Model parameters.** The units Cmmol and gDW refer to mmol carbon and gram dry weight, respectively.

where the hat symbol ($\hat{.}$) denotes the estimated value for a variable or parameter. The value $\hat{g}$ corresponds to 3.25% of the total biomass, close to previous estimates [4, 11].

Metabolic fluxes were derived from data by [5] including the glucose uptake rate $v_{mc}$ [$\text{mmol}_{\text{glc}}$ gDW$^{-1}$ h$^{-1}$], the acetate secretion rate $v_{mef}$ [$\text{mmol}_{\text{ace}}$ gDW$^{-1}$ h$^{-1}$], and the growth rate $\mu$ [h$^{-1}$]. The measured fluxes, together with the degradation rate and the total biomass density, fix the biosynthesis fluxes in the model. In particular, for glycogen synthesis we obtain from Eq. (S2) at steady state:

$$\hat{v}_g = (\hat{\mu} + \hat{\gamma})\,\hat{g} = 0.84 \text{ Cmmol gDW}^{-1}\text{ h}^{-1}. \tag{S22}$$

Glycogen synthesis becomes significant when the supply of carbon source exceeds the demand for macromolecular synthesis, that is, when the use of central metabolites for other processes, approach saturation. This involves complex regulatory mechanisms of the affinity and rate of a key enzyme in the glycogen synthesis pathway, ADP-glucose pyrophosphorylase [4, 12, 13]. We approximated these mechanisms here by choosing a high value for the half-saturation constant of the reaction, that is,

$$\hat{K}_g = 10\,\hat{K}_r, \tag{S23}$$

where $\hat{K}_r = 0.29$ Cmmol gDW$^{-1}$ is the half-saturation constant for protein synthesis (Table S1.4). The ATP half-saturation constant has been assumed to be equivalent to that of the other macromolecular synthesis reaction, *i.e.*

$$K_{ag} = K_{ar} = K_{amu} = 0.0009 \text{ mmol gDW}^{-1} \tag{S24}$$

Together with estimations of the glycogen synthesis rate and the concentration of enzymes in central carbon metabolism (Table S1.3), we can now derive a value for the unknown catalytic constants $k_g$. Using Eq. S2, we thus obtained

$$\hat{k}_g = 3.2 \text{ h}^{-1}. \tag{S25}$$

The coefficient $n_g$ describes the ATP costs of glycogen synthesis. Based on [14], for each glucose residue integrated into the glycogen chain, one ATP is consumed for G6P synthesis and one for the polymeration reaction. Assuming an average chain length of 10 glucose units, we can estimate the ATP cost for glycogen synthesis as

$$n_g = 20/60 = 0.33 \text{ mmol ATP /Cmmol} \tag{S26}$$

All parameter values used in the model are summarized in Table S1.4.

| Rates | Unit | Value | Reference |
|---|---|---|---|
| $\hat{\mu}$ | $h^{-1}$ | $0.61 \pm 0.01$ | [5, 6] |
| $\hat{\gamma}$ | $h^{-1}$ | 0.027 | [15, 16] |
| **Uptake, secretion, biosynthesis fluxes** | | | |
| $\hat{v}_{mc}$ | $\mathrm{Cmmol\,gDW^{-1}\,h^{-1}}$ | $49.6 \pm 5$ | [5] |
| $\hat{v}_{mer}$ | $\mathrm{Cmmol\,gDW^{-1}\,h^{-1}}$ | 4.6 | Derived |
| $\hat{v}_{mef}$ | $\mathrm{Cmmol\,gDW^{-1}\,h^{-1}}$ | $9.8 \pm 3.0$ | [5] |
| $\hat{v}_{mu}$ | $\mathrm{Cmmol\,gDW^{-1}\,h^{-1}}$ | 6.5 | Derived |
| $\hat{v}_{r}$ | $\mathrm{Cmmol\,gDW^{-1}\,h^{-1}}$ | 19.2 | Derived |
| $\hat{v}_{g}$ | $\mathrm{Cmmol\,gDW^{-1}\,h^{-1}}$ | 0.8 | Derived |
| **Total biomass concentration** | | | |
| $1/\hat{\beta}$ | $\mathrm{Cmmol\,gDW^{-1}}$ | $40.65 \pm 2.0$ | [17] |
| **Protein concentrations** | | | |
| $\hat{m}_u$ | $\mathrm{Cmmol\,gDW^{-1}}$ | $11.1 \pm 0.5$ | [2, 8, 17] |
| $\hat{r}$ | $\mathrm{Cmmol\,gDW^{-1}}$ | $13.2 \pm 0.6$ | [2, 8, 17] |
| $\hat{m}_c$ | $\mathrm{Cmmol\,gDW^{-1}}$ | $2.7 \pm 0.1$ | [2, 8, 17] |
| $\hat{m}_{er} + \hat{m}_{ef}$ | $\mathrm{Cmmol\,gDW^{-1}}$ | $3.0 \pm 0.1$ | [2, 8, 17] |
| $\hat{m}_{er}$ | $\mathrm{Cmmol\,gDW^{-1}}$ | 1.9 | Derived |
| $\hat{m}_{ef}$ | $\mathrm{Cmmol\,gDW^{-1}}$ | 1.1 | Derived |
| **Metabolite concentrations** | | | |
| $\hat{c}$ | $\mathrm{Cmmol\,gDW^{-1}}$ | $0.35 \pm 0.002$ | [6, 10] |
| $\hat{g}$ | $\mathrm{Cmmol\,gDW^{-1}}$ | $1.32 \pm 0.002$ | [2, 3] |
| $\hat{a}^*$ | $\mathrm{mmol\,gDW^{-1}}$ | $0.009 \pm 0.0002$ | [6] |
| $\hat{a}$ | $\mathrm{mmol\,gDW^{-1}}$ | $0.011 \pm 0.0006$ | [6] |
| $\hat{a}_0$ | $\mathrm{mmol\,gDW^{-1}}$ | $0.020 \pm 0.0008$ | [6] |
| **Concentration of other biomass** | | | |
| $\hat{u}$ | $\mathrm{Cmmol\,gDW^{-1}}$ | 8.9 | Derived |

**Table S1.3: Growth and degradation rates, uptake, secretion and biosynthesis fluxes, and protein and metabolite concentrations from published data sets for the case of batch growth of *E. coli* in minimal medium with glucose.** Details on the conversion and use of the reported values can be found in the description of the original model [1].

| Parameter | Glucose | Unit |
|---|---|---|
| $\hat{\rho}_{ru}$ | 1.17 | - |
| $\hat{\rho}_{mef}$ | 1.5 | - |
| $\hat{k}_r$ | 2.9 | $h^{-1}$ |
| $\hat{k}_{mu}$ | 1.03 | $h^{-1}$ |
| $\hat{e}_s$ | 18.3 | $h^{-1}$ |
| $\hat{k}_{mer}$ | 5.0 | $h^{-1}$ |
| $\hat{k}_{mef}$ | 17.4 | $h^{-1}$ |
| $\hat{k}_a$ | 2340 | $h^{-1}$ |
| $\hat{k}_g$ | 3.2 | $h^{-1}$ |
| $\hat{K}_r$ | 0.29 | $Cmmol\,gDW^{-1}$ |
| $\hat{K}_{mu}$ | 0.29 | $Cmmol\,gDW^{-1}$ |
| $\hat{K}_{mer}$ | 0.29 | $Cmmol\,gDW^{-1}$ |
| $\hat{K}_{mef}$ | 0.29 | $Cmmol\,gDW^{-1}$ |
| $\hat{K}_g$ | 2.9 | $Cmmol\,gDW^{-1}$ |
| $\hat{K}_{ar}$ | 0.0009 | $mmol\,gDW^{-1}$ |
| $\hat{K}_{amer}$ | 0.0011 | $mmol\,gDW^{-1}$ |
| $\hat{K}_{amef}$ | 0.0011 | $mmol\,gDW^{-1}$ |
| $\hat{K}_{amu}$ | 0.0009 | $mmol\,gDW^{-1}$ |
| $\hat{K}_{ag}$ | 0.0009 | $mmol\,gDW^{-1}$ |
| $\hat{n}_{mer}$ | 4.3 | $mmol\,Cmmol^{-1}$ |
| $\hat{n}_{mef}$ | 2.0 | $mmol\,Cmmol^{-1}$ |
| $\hat{n}_r$ | 0.77 | $mmol\,Cmmol^{-1}$ |
| $\hat{n}_{mu}$ | 0.65 | $mmol\,Cmmol^{-1}$ |
| $\hat{n}_g$ | 0.33 | $mmol\,Cmmol^{-1}$ |

**Table S1.4: Values of the kinetic parameters in the model, in the case of batch growth of *E. coli* in minimal medium with glucose.** The values were derived by means of the calibration procedure of the previous model [1], modified for glycogen synthesis in this work.

#### Code for model calibration and simulation

In order to produce the plots with rate-yield phenotypes (Figures 5-6), we uniformly sampled combinations of resource allocation parameters $x_r$, $x_c$, $x_{er}$, and $x_{ef}$ in such a way that their sum equals 1-$x_u$, where $x_u$ was sampled from a reduced interval determined from the data [1, 8]. Starting from random initial conditions, the system was simulated for each combination of resource allocation parameters until a steady state was reached, where rate and yield were computed from the fluxes and concentrations at steady state.

Whereas every strategy gives rise to a unique rate-yield phenotype, the inverse is not true: different strategies can produce the same growth rate and growth yield. The set of resource allocation parameters corresponding to the wild-type phenotype $(\mu^*, Y^*)$ (Fig. 6C in the main text) was determined using an optimization approach, under the assumption $x_u = 0.37$ (measured value, [1, 8]). Specifically, we searched the $x$ vectors minimizing the difference $\mu - \mu^*$, under the constraints that $Y = Y^*$ and $\sum x_i \leq 1$. Optimization was carried out by means of the `fmincon` function in Matlab R2025.

The model and Matlab simulation code used for generating all figures in the paper are available at `https://gitlab.inria.fr/baldazzi/glycogen-model`. The code includes functions for inferring parameter values from input data on concentrations and fluxes (model calibration), predicting the rate-yield phenotype for a given resource allocation strategy (model simulation), inferring sets of resource allocation strategies capable of producing a given rate-yield phenotype, and analyzing the literature datasets for *E. coli* and *T. lutea*.

[3] G. Eydallin, A. Viale, M. Morán-Zorzano, F. Muñoz, M. Montero, E. Baroja-Fernández, and J. Pozueta-Romero, "Genome-wide screen-

ing of genes affecting glycogen metabolism in *Escherichia coli* K-12," FEBS Lett., vol. 581, no. 16, pp. 2947–53, 2007.

[11] M. Schaechter, J. Ingraham, and F. Neidhardt, Microbe. Washington, DC: ASM Press, 2006.
